# Mouse Forebrain Region-specific Molecular Characteristics and Cholinergic Neurons Subtyping: An Integrated Analysis Based on Spatial Multi-omics

**DOI:** 10.1101/2024.12.12.628119

**Authors:** Yujie Chen, Mengyao Han, Jiayue Meng, Dongjian Cao, Nianhao Cheng, Qingming Luo, Jie Yang, Guoqing Zhang

**Affiliations:** Bio-Med Big Data Center, Chinese Academy of Sciences Key Laboratory of Computational Biology, Shanghai Institute of Nutrition and Health, University of the Chinese Academy of Sciences, Chinese Academy of Sciences, Shanghai 200031, China; Putuo People’s Hospital, Shanghai Key Laboratory of Signaling and Disease Research, School of Life Sciences and Technology, Tongji University, Shanghai 200092, China; Britton Chance Center and MoE Key Laboratory for Biomedical Photonics, Wuhan National Laboratory for Optoelectronics-Huazhong University of Science and Technology, Wuhan 430074, China; State Key Laboratory of Digital Medical Engineering, Key Laboratory of Biomedical Engineering of Hainan Province, School of Biomedical Engineering, Hainan University, Haikou 570228, China

**Keywords:** Multi-omics, Spatial transcriptome-proteome, Mouse forebrain region, Cholinergic neurons-celltyping, RCA-FISH

## Abstract

The forebrain regions displays distinct yet under-characterized gene expression patterns. This study analyzed region-specific feature genes in forebrain regions and Isocortex sub-regions at both transcriptomic and proteomic levels. The key finding is the observation of low correlation but high functional similarity between mRNA and protein expression, which offers insights into the connection between gene expression forms in neuronal pathways and the neuronal activity state. Cholinergic neurons (CNs), play a vital role for the forebrain’s sensory and motor regulation. With Spatial-transcriptome and immunofluorescence joint analysis (STIF), which overcomes the limited resolution of the 10X Visium system, CNs- and CN-subtypes specific feature genes in the striatum and basal forebrain were identified, which providing crucial insights into the heterogeneity and functional diversity of these neuronal populations. The spatial distribution and expression pattern of the feature genes in this study were validated with either external datasets or rolling circle amplification-FISH, coincident results were revealed.

## Introduction

The brain is a complex organ, plays a crucial role in cognition and disease, the spatial organization of the brain is fundamentally related to its function (1, 2). The forebrain is the seat of higher order brain functions, different brain regions show distinct gene expression patterns (3, 4). Comprehensive understanding of intricate biological systems of various brain regions requires spatially resolved transcriptomic and proteomic characterization (5, 6). However, the joint analysis strategy of transcriptomic and proteomic in neuroscience is still in its infancy. The existing strategies for generating high-throughput spatial expression profiles were developed primarily on the spatial transcriptome, which enabled the simultaneous measurement of morphological features and transcriptional profiles of the same cells or regions in tissues (7–9). Recently developed imaging-based spatial transcriptomics technologies, such as 10x Genomics Xenium and Vizgen MERSCOPE MERFISH, offer single-cell resolution. However, their current limitation is that they can only measure 400 or 1000 targeted transcripts. To systematically investigate the molecular features of the major forebrain regions, it is essential to capture a sufficient number of untargeted genes with in situ technology. Consequently, the 10X Visium platform was utilized for this purpose.

Proteins are the main functional components in all tissues, but the development of spatial proteomics was still in the initial stage. Most spatial proteomics were based on targeted antibodies (10, 11). The number of detected proteins were limited from several to dozens, which is inadequate for the analyzing molecular features and difficult to be parallelly analyzed with the high-throughput spatial transcriptome data. In order to detect a sufficient number of untargeted proteins in brain regions, we performed mass spectrometry (MS)-based spatial proteomics with laser capture microdissection (12). Subsequently, we conducted a comprehensive joint analysis by integrating spatial transcriptome and proteome data to explore the association between mRNAs and proteins in brain regions and neuronal pathways.

Cholinergic neurons (CNs) in the forebrain play essential roles in regulating sensory and motor functions as well as cognitive behaviors. Two main types of CNs are cholinergic interneurons (CINs) in the striatum and cholinergic projection neurons in the basal forebrain (BFCNs) (13). Previous studies found CINs exhibit substantial diversity in their physiology, morphology and connectivity (14), and the projections of BFCNs to the forebrain and midbrain showed distinct specificity which suggested the existence of distinct subtypes of CINs and BFCNs that serve different regulatory functions in the brain (15, 16). However, the molecular subtype classification of these cells is still incomplete, primarily due to the current limitations of cell typing research strategies.

Flow cytometry and single-cell RNA sequencing technology are commonly used in neuronal typing research. However, only a limited number of CNs could be detected in vitro (17), which makes it challenging to investigate their subtypes. To obtain a sufficient amount of transcriptional data from hundreds of CINs and BFCNs, we utilized the 10X Visium Spatial to capture the transcripts of the striatum and basal forebrain across multiple brain sections. But the spot size of this system is 55 µm and each spot still contains around 7 cells for the striatum and basal forebrain. To minimize the interference from the surrounding cells, we introduced a novel strategy called "STIF", which integrated 10X transcriptome data with immunofluorescence (IF) to identify distinct features and subtypes of CINs and BFCNs.

In this article, we have provided easily implementable strategies for conducting integrative analysis of multi-omics data and extracting cellular features from cell mixed environments. With the joint approach, we identified 29 and 16 molecular features in the forebrain regions and Isocortex sub-regions. Of these, 18 were brain region feature genes and 16 were sub-region feature genes newly found in our study. These feature genes were closely linked to brain region functions and validated using external datasets, which revealed similar expression pattern in terms of expression and spatial distribution. Moreover, the discovery of a low correlation but high functional similarity between mRNA and protein expression patterns in brain regions has provided insights into the diverse gene expression forms (mRNA or protein) within neuronal pathways. This divergence in expression may be associated with the state of neuronal activity. With the implementation of STIF, according to the expression specificity, we successfully extracted 12 feature genes of CINs and BFCNs from the cell-blended data. These feature genes are closely linked to the functions of CNs, 6 of which were newly discovered in our study. To validate these genes, we reanalyzed Munoz-Manchado’s scRNA-seq dataset and found that most of them were also specifically expressed in Chat-expressing cells. Moreover, with the HEGs of CNs, 4 subtypes of striatal cholinergic interneurons and 3 subtypes of basal forebrain cholinergic projection neurons were categorized, providing crucial insights into the heterogeneity and functional diversity of these neuronal populations. Their feature genes have been validated through rolling circle amplification-FISH (RCA-FISH). The expression pattern of these feature genes were greatly coincident between the 10X Visium and RCA-FISH results.

## Results

### Achievement and registration of spatial multi-omics data

The forebrain, which plays a crucial role in higher-order brain functions, exhibits unique gene expression patterns that have not been fully characterized at both the transcriptomic and proteomic levels (2). To simultaneously explore the molecular features at both levels, we obtained 8 groups of immediately adjacent coronal sections (range from Bregma 0.74 ∼ -0.48) from the mouse forebrain using cryosection **(Figure 1A)**.

**Figure 1.**
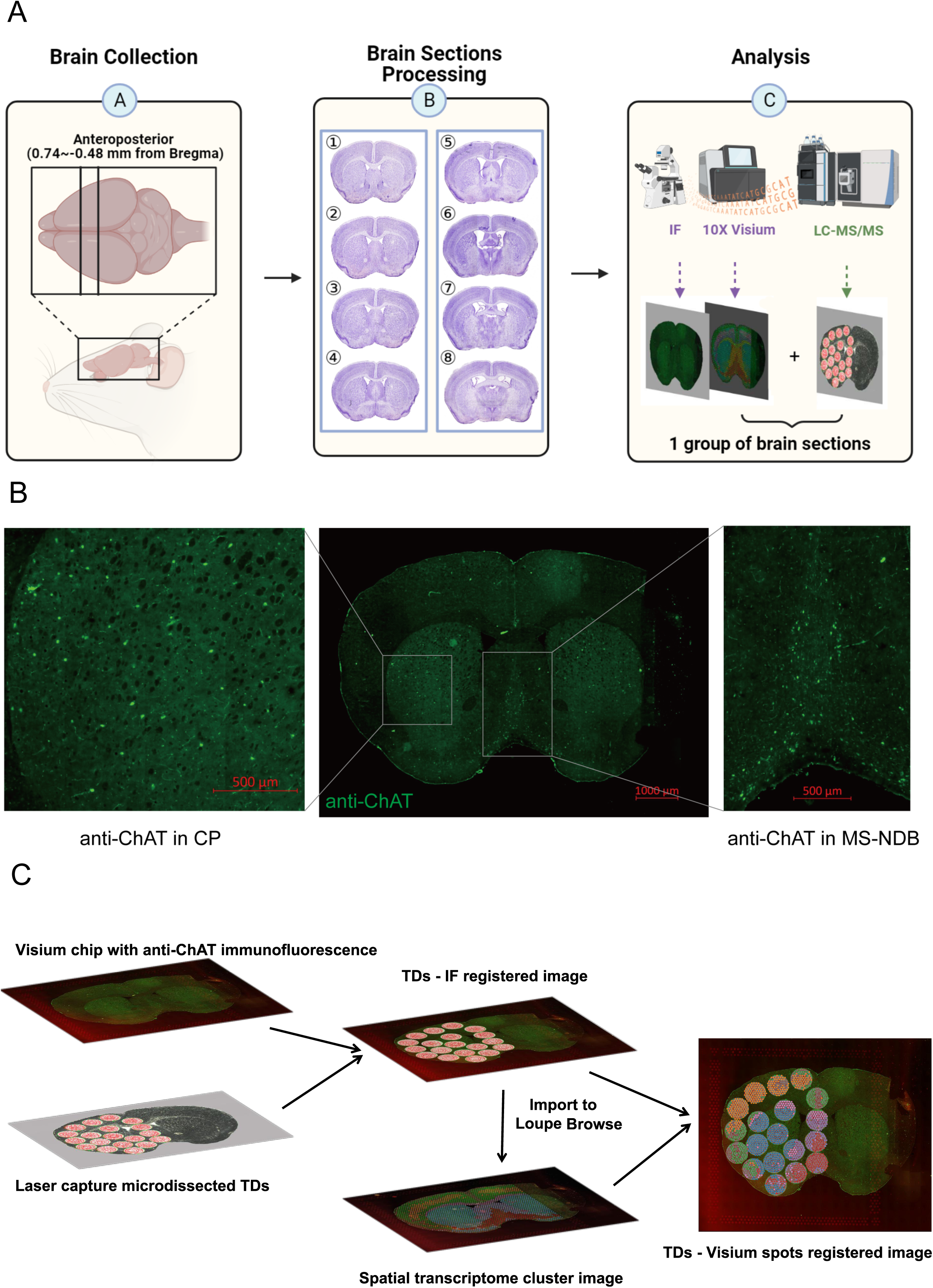
Multi-omics data registration of mouse forebrain. **A**. The strategy to achieve spatial multi-omics data with the adjacent sections of the mouse forebrain. 8 groups of immediately adjacent coronal sections (Bregma 0.74∼-0.48) were sampled at intervals. 10μm section was collected for in situ 2D-RNAseq of spatial transcriptome and anti-ChAT immunostaining (n=8), 20μm immediately adjacent section was collected for spatial proteome (n=8). **B.** Anti-ChAT IF of the mouse forebrain section. The antibody labeled CNs, located in the striatum (left) and the basal forebrain (right). **C.** The workflow of registering spatial transcriptome and spatial proteome data. The brain section on Visium chip with anti-ChAT IF was first registered with the immediately adjacent microdissected brain section. The TDs – IF registered image was imported to the Loupe Browser to associate with the Visium spots. The section contained IF signal, transcriptome and proteome information.

Within each of these 8 groups, a pair of adjacent sections was processed. One of these sections was cryosectioned to a thickness of 10µm, followed by hybridization onto a spatially barcoded array for the collection of in situ 2D-RNAseq data. Before tissue permeabilization, an anti-ChAT antibody was applied to this section. Simultaneously, the immediately adjacent section, with a thickness of 20µm was cryosectioned. To ensure the homogeneity of the sample, the same size tissue domains (TDs) were carefully dissected from these sections using laser capture microdissection and subsequently subjected to analysis through liquid chromatography-tandem mass spectrometry (HPLC-MS/MS) (Figure 1A; **Figure 1B**).

The brain section on Visium chip provided spatial transcriptomic data, while the adjacent microdissected section containing the TDs offered spatial proteomic information. The precise locations of the TDs were documented during microdissection by referencing the cutting marks using bright-field imaging. To associate anti-ChAT signal, spatial transcriptome and proteome data and analyze them synchronously, the brain section on Visium chip with anti-ChAT IF was first registered with the immediately adjacent microdissected brain section according to the outline of the brain sections. The microdissected-Visium chip registered image was then imported to the Loupe Browser according to the fiducial frames, where the Visium spots were positioned. By this way, TDs on microdissected brain section were associated with the Visium spots and anti-ChAT signal on Visium chip (**Figure 1C**). The proteome TDs have an average diameter of 936μm and a thickness of 20µm, which was determined as the threshold for reliable identification of ChAT (Figure S1), corresponds to 75-85 Visium spots with a diameter of 55 μm. The majority of ChAT expressed TDs correspond to the anti-ChAT IF signal, indicating a strong consistency between nearby brain sections.

Thus, this approach allowed us to simultaneously investigate the gene expression patterns at both the transcriptomic and proteomic levels, providing a more comprehensive understanding of the molecular landscape in the forebrain.

### Expression of the in-situ mRNA and protein in the major forebrain regions

To ensure precise identification and characterization of the expression patterns within the forebrain regions, accurate delineation of brain regions is essential. We utilized the Allen Brain Atlas as a template to delineate brain regions and made adjustments to the delineation based on patterns observed in the anti-ChAT signal and spatial transcriptome clustering (Figure S2A). Only the TDs that do not span multiple brain regions can be utilized to analyze the characteristics of a specific brain region. Therefore, we combined these smaller brain regions into their respective parent structural levels: Isocortex, OLF-PIR (Olfactory-Piriform area), STR-CP&ACB (Striatum, Caudoputamen & Nucleus accumbens), PALm&PALv (Pallidum, medial region & ventral region) and Hypothalamus (**Figure 2A**; Figure S2A).

**Figure 2.**
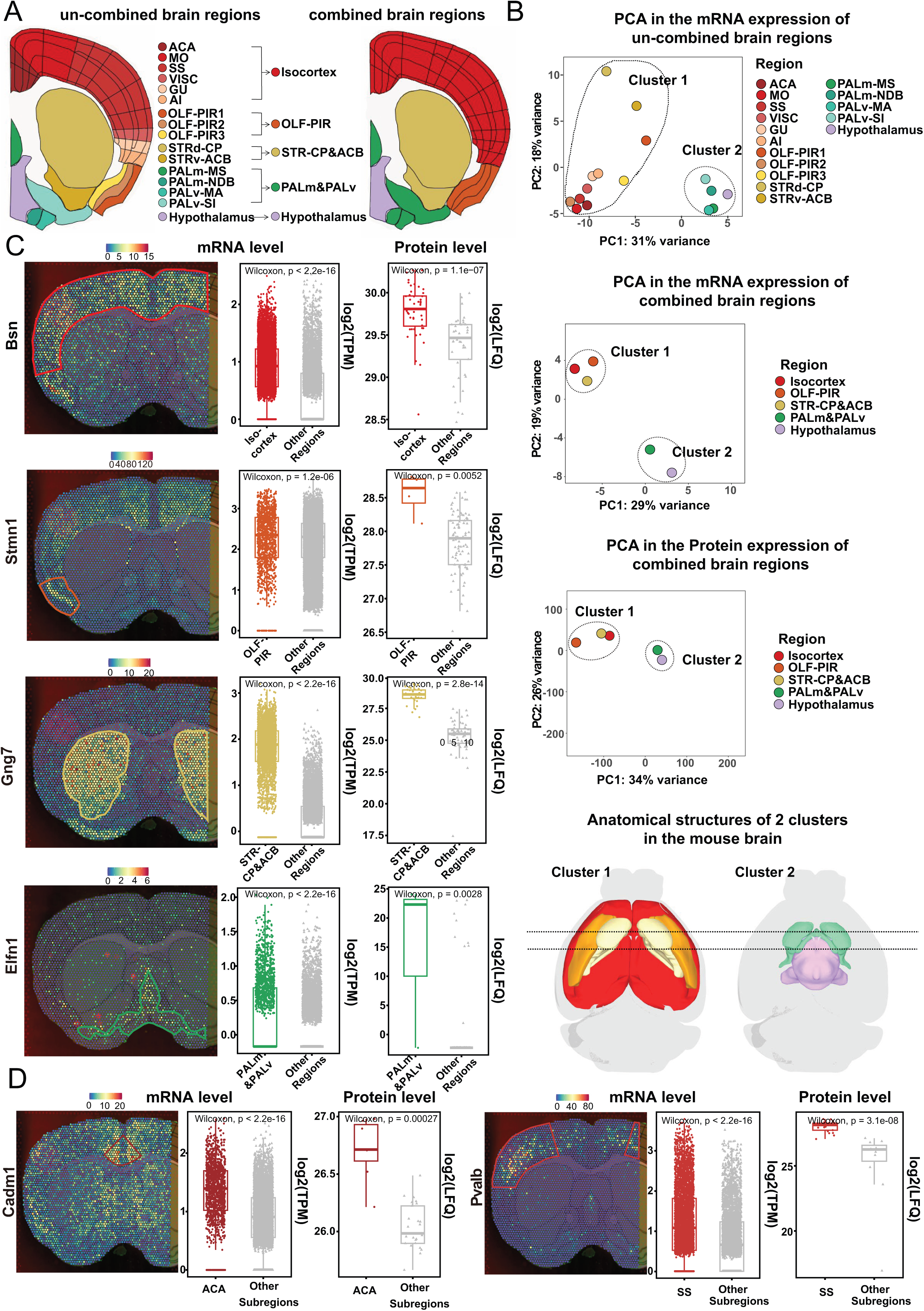
Analysis of the in situ mRNA and protein expression in brain regions. **A**. The relationship of uncombined and combined brain regions. The Isocortex was combined from anterior cingulate area (ACA), somatomotor areas (MO), somatosensory areas (SS), visceral area (VISC), gustatory areas (GU), agranular insular area (AI). The OLF-PIR was combined from piriform area-molecular layer (OLF-PIR1), piriform area-pyramidal layer (OLF-PIR2) and piriform area-polymorph layer (OLF-PIR3).The STR-CP&ACB was combined from striatum dorsal region-caudoputamen (STRd-CP) and striatum ventral region-nucleus accumbens (STRv-ACB). The PALm&PALv was combined from pallidum-medial region-medial septal nucleus (PALm-MS), diagonal band nucleus (PALm-NDB), pallidum-ventral region-substantia innominata (PALv-SI) and magnocellular nucleus (PALv-MA). **B.** PCA and anatomical distribution of brain regions. PCA of uncombined and combined brain regions according to mRNA expression; PCA of combined brain regions according to protein expression; anatomical distribution of the brain region clusters in the mouse brain. **C.** The mRNA spatial distribution of Isocortex feature gene Cd34, OLF-PIR feature gene Stmn1, STR-CP&ACB feature gene Nexn and PALm&PALv feature gene Elfn1 (left). The boxplot of the mRNA expression and protein abundance of these genes in relevant brain regions (Isocoretx, TDs=48; OLF-PIR, TDs=4; STR-CP&ACB, TDs=30; PALm&PALv, TDs=3; Hypothalamus, TDs=4; wilcoxon test) (right). **D.** The mRNA spatial distribution of ACA feature gene Cadm1 and SS feature gene Pvalb (left). The boxplot of the mRNA expression and protein abundance of Cadm1 and Pvalb in the Isocortex (ACA, TDs=7; SS, TDs=21; wilcoxon test) (right).

To further explore the relationship between expression and anatomical structures, we performed principal component analysis (PCA) for brain regions at both transcriptome and proteome levels. Two clusters diverged from each other (**Figure 2B**). Cluster 1 included the Isocortex, STR-CP&ACB and OLF-PIR, which were distributed in the peripheral region of the forebrain. Cluster 2 included the PALm&PALv and the Hypothalamus, which were located in the central region of the forebrain (Figure 2B). Thus, the distance of the PCA clusters was related to the anatomical distribution of the brain regions in both transcriptome and proteome analysis.

For each region, we obtained highly expressed genes (HEGs) and highly expressed proteins (HEPs) by comparing transcriptome and proteome expression with other brain regions respectively (Figure S2B). The top 20 HEGs and HEPs were screened based on log2-fold-change in the 5 brain regions and were showed in Figure S2C. To ascertain the reliable transcriptomic and proteomic features, we compared the total HEGs and HEPs in all 5 brain regions (Table S1), more than 200 overlapped genes were achieved. Their spatial distribution was examined at both mRNA and protein expression level. Among them, 29 genes with brain regions specificity were filtered out as molecular features (**Figure 2C**; Table S1; Figure S2D). We examined the distribution of these genes in the literature. Among them, 18 genes were not reported to exhibit high expression at both mRNA and protein levels in their respective brain regions previously. Remarkably, we performed sub-regions molecular feature analysis in the Isocortex for the first time. According to the expression and region-specificity of mRNA and protein, we achieved 3 molecular features for anterior cingulate area (ACA) and 13 molecular features for somatosensory areas (SS) (**Figure 2D**; Figure S3A).

The Isocortex and OLF-PIR belong to the cerebral cortex. The cerebral cortex is the seat of cognitive abilities and composed of an extraordinary number of cells (18). In our study, we found some synapse related genes were highly expressed in the cerebral cortex, associated with the regulation of synaptic vesicles or plasticity. For example, *Bsn* plays a role in regulating the synaptic vesicle cycle (19). *Grin2a* and *Snap25* are involved in the regulation of synaptic plasticity (20, 21). Dnm1 encodes a GTPase necessary for vesicle endocytosis in neurons (22). Sh3gl2 contributes to the positive regulation of vesicle scission, while Stx1a is involved in synaptic vesicle exocytosis (23, 24). Additionally, Dlgap1 is responsible for regulating postsynaptic neurotransmitter receptor activity (25).

The ACA and SS are sub-regions of the Isocortex, exhibit distinct functions. The ACA, which is an integral part of the prefrontal cortex in mice, plays a crucial role in supporting cognitive functions such as attentional processes, motion planning and execution, as well as remote memory, fear and pain (26). On the other hand, the SS is primarily responsible for processing sensory information from different parts of the body (27). The majority of the ACA feature genes were found to be associated with the regulation of synaptic vesicles, implying their function in synaptic transmission. While the SS feature genes were primarily related to the neuromuscular junction and mechanical stimulus response, suggesting their role in processing sensory information.

The striatum, as the principal input nucleus of the basal ganglia, plays a crucial role in motor functions, motivation, and reward-related learning (28, 29). Signaling via G protein-coupled neurotransmitter receptors is essential for motor control (30). We found some G protein related genes were highly expressed in the STR-CP&ACB. The majority of the molecular features in the STR-CP&ACB were related to GPCR signaling. For example, Gnal and Gng7 encode G-protein subunits, while Rasd2 and Rem2 function as GTPases (31–34). Rgs9 forms a complex with G-protein subunits (35). Adcy5, Pde10a and Pde1b play crucial roles in controlling the synthesis and hydrolysis of the second messenger cAMP, serving as key points for GPCR signal integration (36–38). Additionally, Gprin3 acts as a G-protein-regulated inducer of neurite outgrowth (39).

GABAergic neuron was the major neuronal type of the PALm&PALv (40). *Elfn1*, highly expressed in PALm&PALv, involved in synapse organizations, localized to axons of the long-range GABAergic neurons (41). The hypothalamus itself contains several types of neurons that release different hormones (42). Oxytocin (*Oxt*), highly expressed in the hypothalamus, was a posterior pituitary hormone and primarily synthesized in magnocellular neurons (43, 44). In summary, the feature genes were closely linked to the brain region functions.

To validate the newly discovered feature genes, we check the gene expression in ISH data of Allen brain institute, and found 21 out of 29 feature genes were also highly expressed in the corresponding brain regions. We analyzed Ortiz’s dataset to further investigate the expression and distribution of these genes, revealing similar expression patterns and spatial distributions. (Figure S2E; Figure S2F; Figure S3B; Figure S3C) (45).

### The relationship of the mRNA expression and protein abundance in forebrain regions

Gene expression, whether in the form of mRNA or protein, is closely linked to the activity of cells. Cells can control the production and levels of specific proteins by regulating the processes of transcription and translation, thereby influencing the function and activity of the cells (46). To investigate the relationship between mRNA expression and protein abundance of HEGs and HEPs, we first computed the Spearman correlation coefficients across 5 forebrain regions: Isocortex, STR-CP&ACB, OLF-PIR, Hypothalamus and PALm&PALv. We revealed a low correlation between mRNA expression and their corresponding protein abundance in the same brain regions (0.36∼0.39, **Figure 3A**). In contrast, the correlation within mRNA expression or protein abundance in the brain regions exhibited higher values (0.62∼0.66 for mRNA expression and 0.89∼0.93 for protein abundance) (Figure S4A). We also analyzed the gene set that make the higher values of the correlation within mRNA expression and protein abundance in the brain regions, found that most of these genes were associated with neuronal functions, as detailed in Table S2.

**Figure 3.**
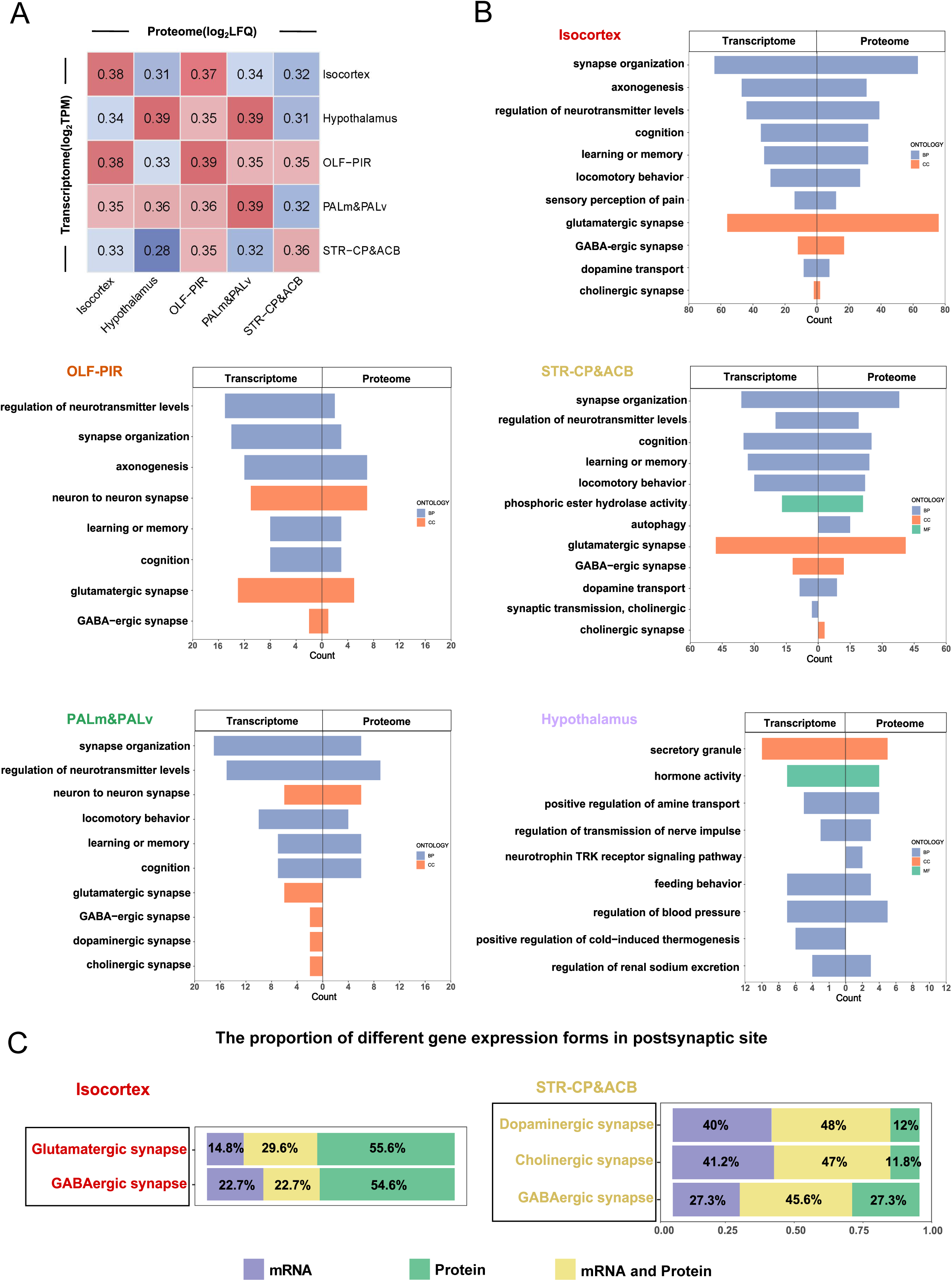
The relationship between the mRNA and protein expression. **A**. Correlation plots of log2 LFQ (proteome) versus log2 TPM values (transcriptome). The color follows the indicated values of correlation coefficient. **B.** Bar plot depicting significantly enriched GO terms associated with HEGs and HEPs in the 5 brain regions. Different colors were used to distinguish Molecular Function, Biological Process and Cellular Component. **C.** The proportion of different expression forms in the postsynapse site (glutamatergic and GABA-ergic synapse in the Isocortex and the dopaminergic, cholinergic and GABA-ergic synapse in the STR-CP&ACB). The bar charts indicated the proportion numbers of the genes which highly expressed mRNAs (purple), highly expressed proteins (green) and highly expressed mRNAs and proteins synchronously (yellow).

We further performed GO enrichment analysis with HEGs and HEPs respectively (Table S2). In the GO analysis, we observed that the enriched functions of HEGs and HEPs were nearly identical (Fisher p-adj: 3.8E-40 ∼ 1.4E-130; Figure S4B). However, the participating genes showed diversity in certain functions, indicating that the expression of mRNAs and proteins may not be synchronized. We analyzed the gene distribution from HEGs and HEPs in the same GO terms, revealing an average overlap ratio of 31.7%. The Isocortex, STR-CP&ACB, OLF-PIR and PALm&PALv were all associated with some common brain functions, such as "neuron to neuron synapse", "synapse organization", "regulation of neurotransmitter levels", "learning or memory", "cognition", "sensory", which corresponded to the original knowledge (47–50). The STR-CP&ACB was enriched into "autophagy" and "phosphoric ester hydrolase activity", which reflected the reported specific function of the striatum (51). The Hypothalamus was reported to be responsible for many neuroendocrine functions, including glucose regulation, thermogenesis, social and sexual behaviors (52). Accordingly, in our results, it was related to "secretory granule", "hormone activity", "regulation of transmission of nerve impulse", "positive regulation of cold-induced thermogenesis", "feeding behavior" (**Figure 3B**).

To clarify the low correlation yet highly similar functions of HEGs and HEPs, we analyzed their profiles across major synaptic pathways in the Isocortex and STR-CP&ACB, which include both presynaptic and postsynaptic sites. We quantified the distribution of HEGs and HEPs at these sites and found that their expression in these pathways was not synchronized (Table S2). Only 22.2-48.0% of the genes exhibited simultaneous high expression in either presynaptic or postsynaptic neurons.

The remaining genes showed high expression exclusively at either the mRNA or protein level. This suggests that the low correlation but highly similar functions of HEGs and HEPs may stem from their involvement in the same pathways, albeit with different expression forms. We further found that the presynaptic site exhibits a higher proportion of HEPs, which may indicate active neurotransmitter production or transport in the Isocortex and STR-CP&ACB (53).

In contrast, the expression forms in postsynaptic site differ between the Isocortex and STR-CP&ACB. In the Isocortex, a significant observation was the increased proportion of HEPs in postsynaptic site, highlighting the primary role of proteins. Conversely, in the STR-CP&ACB, a higher proportion of HEGs in postsynaptic site suggested that mRNAs serve as the primary form of gene expression (**Figure 3C**; Figure S5).

Proteins are the primary components responsible for carrying out cellular functions, and the stability of proteins is crucial for cellular activities (54). Normally, the Isocortex is associated with high synaptic activity (55, 56). An active cortex is necessary for intact cognitive function (57). The higher proportion of HEPs in the postsynaptic site may reflect the active status of the synapse pathways in the Isocortex (Figure 3C).

Previous research has shown that the form in which mRNA exists is closely associated with rapid response. Certain mRNA molecules are present in a stable state within cells, allowing them to quickly translate into proteins upon external stimuli (58). The striatum specified both the contents and structure of behavior, plays a crucial role in motor functions (59). While motor control by striatal neurons was a rapid process, which was regulated in seconds or sub-seconds timescale (29, 60). The higher proportion of HEGs in the postsynaptic site may indicate the rapid response and accurate motor control of the synapse pathways in the STR-CP&ACB (Figure 3C).

### Molecular features identification in the cell-blended forebrain cholinergic system with STIF

Single-cell RNA sequencing recently revealed the transcriptome profiles of CNs, yet their in-situ information remains elusive. In our study, we observed that the spatial transcriptome and proteome data for STR-CP&ACB and PALm&PALv were not distinctly enriched in the cholinergic synapse category in GO analysis. It might be due to the low proportion of the CNs in STR-CP&ACB and PALm&PALv (about 1-3% of all striatal cells and 20% of all basal forebrain cells) (61, 62). The molecular characteristics could be obscured by neurons with a high proportion such as GABA-ergic and glutamatergic neurons. Thus, it is difficult to explore the molecular features of the CNs with our spatial omics data without processing. Each Visium spot contains an average of 7 cells according to hematoxylin-eosin (HE) staining (Figure S6A). Following the anti-ChAT immunostaining, we observed that the CNs were predominantly distributed with only one cell in each spot. To get rid of the interference of the surrounding cells, we integrated immunostaining with spatial transcriptome data and developed a novel method called spatial-transcriptome and immunofluorescence joint analysis (STIF) to uncover the in situ molecular features of these CNs (**Figure 4A**; **Figure 4B**).

**Figure 4.**
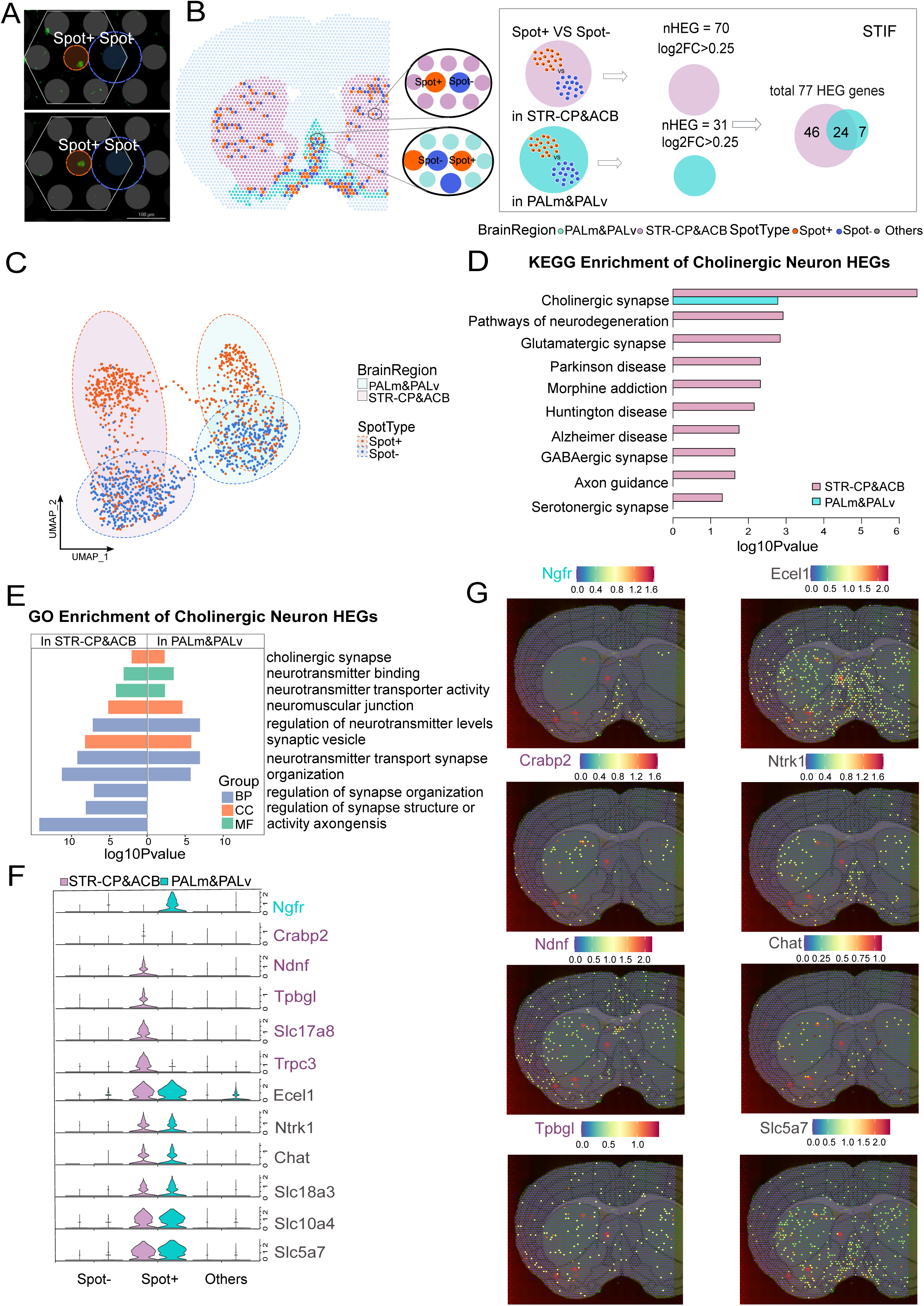
Explore the molecular features of the stratial CINs and BFCNs via STIF. **A**. The definition and selection of the "Spot+" and "Spot-". The 10X Visium spots containing anti-ChAT IF signals (orange circle) were selected to be "Spot+". The spots immediately adjacent to the "Spot+"(within the hexagon), in and around which (blue circle) did not contain any anti-ChAT IF signal were selected to be the "Spot-". **B.** The diagram of STIF workflow. The transcriptomics profiles between "Spot+" and "Spot-" in the STR-CP&ACB (n=408; 402) and the PALm&PALv (n=295; 268) were compared respectively with wilcox test, FDR<0.05, log2FC>0.25. 77 HEGs were identified. **C.** UMAP plot of the "Spot+" and "Spot-" in STR-CP&ACB (n=408; 402) and PALm&PALv (n=295; 268) with 77 HEGs. **D.** Bar plot depicting significantly enriched KEGG pathways associated with HEGs, FDR<0.05. **E.** Bar plot depicting significantly enriched GO terms associated with HEGs in STR-CP&ACB and PALm&PALv, with a threshold of FDR<0.05. **F.** Violin plot of 12 CNs feature genes for "Spot+", "Spot-", "Others" in the STR-CP&ACB (n=408; 402; 5739) and PALm&PALv (n =295; 268; 1514). **G.** The spatial distribution of newly identified CNs feature genes (Ecel1, Crabp2, Ntrk1, Ndnf, Tpbgl and Slc5a7) on the brain section.

We obtained 77 HEGs of the CNs in STR-CP&ACB and PALm&PALv (70 for striatal CINs, 31 for BFCNs) by comparing the transcriptomics profiles between the "Spot+" and "Spot-" (Figure 4B; Table S3). With these 77 genes, we were able to distinctly separate the "Spot+" and "Spot-" in both regions (**Figure 4C**). The well-separated "Spot+" were utilized for subsequent subtyping analysis.

With another comparing strategy, 130 up regulated genes of BFCNs and CINs were identified (Figure S6B). Nevertheless, the clustering result of the "Spot+" and "Spot-" with those genes was farraginous (Figure S6C). Clustering with the top 2000 highly variable genes revealed significant overlap of the "Spot+" and "Spot-" (Figure S6C). Moreover, ARI scores indicated that the STIF strategy, which relied on the 77 genes, more effectively distinguished the "Spot+" and the "Spot-" (Figure S6D). Therefore, accurate extraction of HEGs is crucial for the subsequent analysis.

We performed KEGG enrichment analysis with HEGs in BFCNs and striatal CINs respectively. These genes were observed to be enriched in several pathways associated with neurons and neuronal diseases. In particular, the HEGs of striatal CINs and BFCNs were both enriched in the "cholinergic synapse" pathway (**Figure 4D**). Furthermore, the genes found in striatal CINs were enriched in various brain diseases, including Alzheimer’s disease, Parkinson’s disease and Huntington’s disease. These diseases have been reported to be correlated with CINs in the straitum (63, 64), confirming the effectiveness of "STIF". We also performed GO enrichment analysis, found the HEGs of striatal CINs and BFCNs were both enriched in "cholinergic synapse", "neurotransmitter transport" and "synapse organization", which further suggests the validity of "STIF" (**Figure 4E**).

To identify the molecular features for the striatal CINs and BFCNs, we further screened these HEGs by the expression specificity and their spatial distributions on brain sections. 12 genes were identified as the features of the CNs (**Figure 4F**; **Figure 4G**; Figure S6E; Figure S6F). *Chat*, *Slc5a7*, *Slc18a3*, *Ntrk1*, *Slc10a4* and *Ecel1* were found to be expressed in both the striatal CINs and BFCNs. *Trpc3*, *Slc17a8*, *Tpbgl*, *Crabp2* and *Ndnf* were specific to CINs, while *Ngfr* was specific to BFCNs (Figure 4F; Figure S6E). Among them, *Ecel1*, *Crabp2*, *Ndnf*, *Tpbgl*, *Slc5a7* and *Ntrk1* were newly identified as being highly expressed in mouse’s CINs or BFCNs in our study (Figure 4F; Table S3).

*Chat, Slc5a7, Slc18a3, Ntrk1* and *Slc10a4* were demonstrated to be expressed in CNs, the majority of them were enzymes and transporters that regulate cholinergic neuronal function (Table S3). *Ecel1*was found to be involved in nerve regeneration events (65). In striatum, *Trpc3, Slc17a8* and *Crabp2* were demonstrated to be expressed in striatal CINs, ensured the normal transmission of neural signals (Table S3). *Tpbgl* was reported to express in mouse’s basal ganglia and Ndnf was a neurotrophic factor (66, 67). *Ngfr* (p75NTR), a neurotrophin receptor, which mediated BFCNs neuronal death. Knockout of p75NTR leaded to an increase in the number of BFCNs and also affected the size of BFCNs (68). In summary, most of the molecular feature genes were closely linked to the CNs functions.

To validate the newly discovered feature genes, we utilized Munoz-Manchado’s single-cell RNA sequencing (scRNA-seq) dataset to reanalyze and found that the majority of these genes were also specifically expressed in the Chat-expressing cells (Figure S6G) (17).

### Subtype identification and spatial distribution of CINs and BFCNs in forebrain

The striatum is commonly divided into dorsal and ventral striatum (DS, VS) (14). In rodents, the DS often referred to CP and the VS to ACB. The CP consists of two functionally distinct regions: the dorsomedial striatum (DMS) mediating goal-directed actions, cognition and behavior; the dorsolateral striatum (DLS) mediating habitual actions and motor control (69, 70). The boundary of two subregions could be indicated by *Cnr1* or *Crym* (**Figure 5A**; Figure S7A).

**Figure 5.**
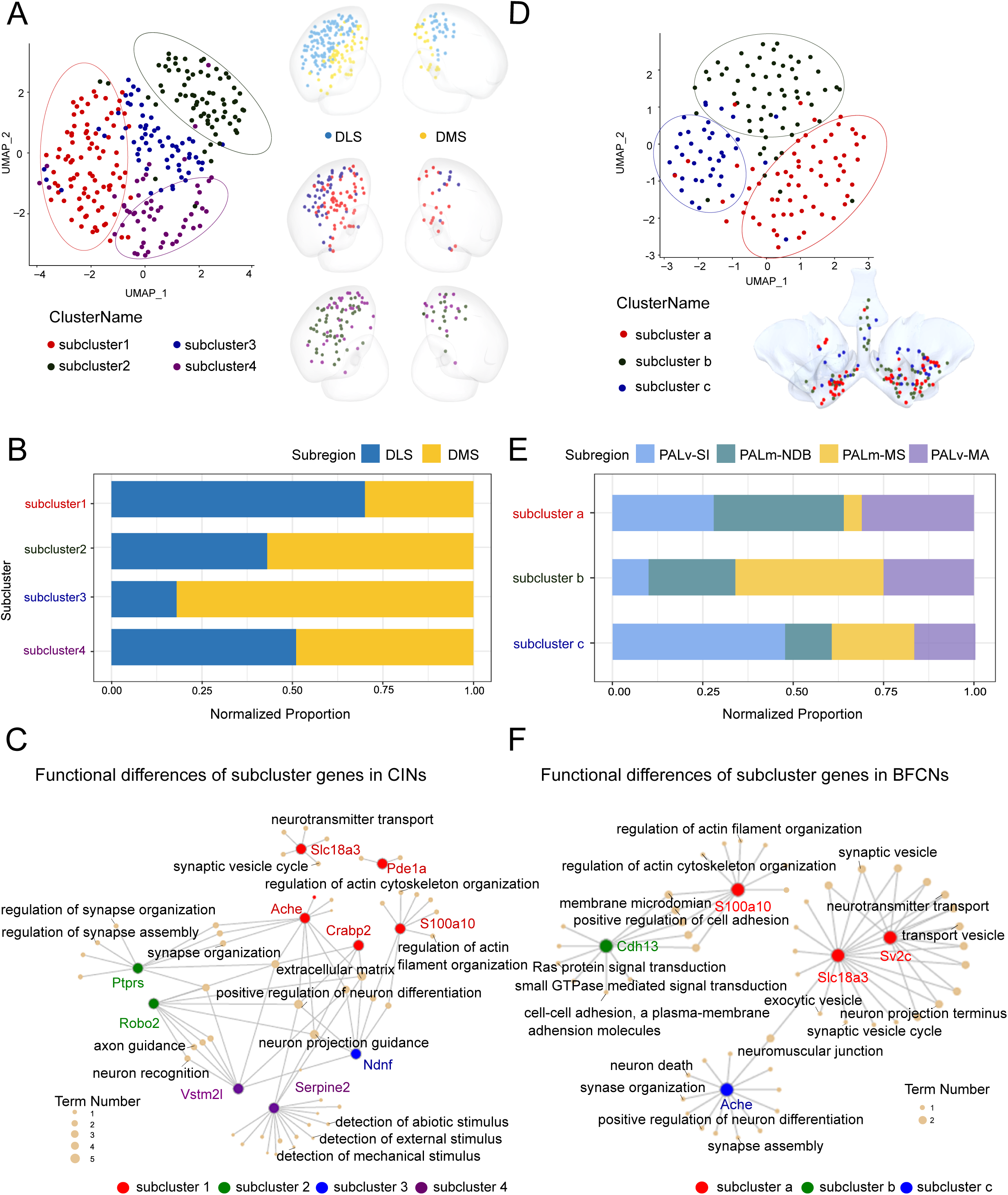
Subtype identification of striatal CINs and BFCNs. **A.** Left, UMAP plot of 4 subclusters for STR CINs "Spot+" (subcluster1=88; subcluster2=70; subcluster3=61; subcluster4=55). Right, 3D spatial visualization show the distribution of DLS (blue) and DMS (yellow) at the top, the subclusters distribution of CINs at the bottom. The boundary of DLS and DMS were determined according to the expression of Cnr1 and Crym. **B.** The distribution proportions of striatal CIN subclusters in DLS and DMS. **C.** The GO enrichment of subcluster feature genes in CINs. **D.** Left, UMAP plot of 3 subclusters for BFCNs "Spot+" (subcluster a=63; subcluster b=59; subcluster c=39). Right, 3D spatial visualization of the BFCNs subclusters distribution in PALm&PALv. **E.** The distribution proportions of BFCN subclusters in MA, MS, NDB and SI. **F.** The GO enrichment of subcluster feature genes in BFCNs.

CINs were previously reported to have two distinct morphological populations in the CP (14), indicating the existence of potential sub-clusters. To identify potential subclusters in situ, well-separated "Spot+" in STR-CP&ACB (n=274) were clustered with CINs HEGs, resulting in the identification of 4 subclusters (Figure 5A). Interestingly, we found their spatial distributions had bias to DLS or DMS. Subclusters 2 and 4 were distributed across both the DMS and DLS, whereas subcluster 1 was predominantly localized to the DMS, and subcluster 3 was primarily confined to the DLS (Figure 5A; **Figure 5B**; Figure S7B). HEGs were identified in these subclusters respectively (Table S3, Figure S7D). We conducted a clustering analysis of the enriched GO terms to investigate the functional differences among the feature genes of CIN subclusters. Subcluster 1 was primarily associated with GO terms such as the ’synaptic vesicle cycle’ and ’regulation of actin cytoskeleton organization’. Subcluster 2 showed significant enrichment in terms related to ’regulation of synapse organization’. Subcluster 3 was linked to GO terms involved in ’neuron projection guidance’, while Subcluster 4 was associated with ’detection of mechanical stimulus’(**Figure 5C**). These findings indicated the correlation between spatial distribution and functional diversity of CINs.

BFCNs historically categorized into Ch1, Ch2, Ch3 and Ch4 groups based on their anatomical locations: Ch1 (Medial septal nucleus - MS), Ch2 (Diagonal band nucleus - NDB), Ch3 (Magnocellular nucleus - MA), and Ch4 (Substantia innominata - SI) (71). In our studies, to explore the subclusters of BFCNs at molecular level, well-separated "Spot+" in PALm&PALv (n=161) were clustered with BFCNs HEGs and acquired 3 subclusters (**Figure 5D**). Interestingly, their distributions differed from the anatomical classification and exhibited a mixed state among all the PALm&PALv regions. Subcluster b was predominantly found in MS-NDB but less so in SI, subcluster a was more prevalent in NDB, whereas subcluster c was more concentrated in SI (Figure 5D; **Figure 5E**; Figure S7C). Previously, BFCNs in MS-NDB were identified to have 3 subpopulations with distinct projection specificity and their distribution were also mixed (16), which indicated that different subpopulations of BFCNs may be different in both transcriptome and projection profiles. In these 3 BFCNs subclusters, we identified several HEGs (Table S3, Figure S7E). Clustering analysis of enriched GO terms revealed distinct functional profiles among BFCN subclusters. Subcluster a was primarily associated with GO terms related to ’neurotransmitter transport’ and ’regulation of actin cytoskeleton organization’. Subcluster b showed significant enrichment in terms related to cell adhesion and ’small GTPase mediated signal transduction’, while subcluster c was linked to terms involved in ’synapse organization’ (**Figure 5F**).

### Validation via rolling circle amplification-FISH (RCA-FISH) for the subtypes feature genes

To verify the feature genes of the subtypes in CINs and BFCNs respectively, we performed the RCA-FISH experiment on the forebrain sections. We noticed that the anti-ChAT signals were highly overlapped with the signals of Chat-RCA-FISH, therefore both could be used as the marker of the CNs (Figure S8A). We quantified the RCA-FISH signals of these subtype features genes in each Chat expressed cell and defined it as the expression level. We also quantified the proportions of single-positive (Chat-expressed) and double-positive (co-expressed with a feature gene) signals in striatal CINs and BFCNs, finding that over 55% of the signals in these neurons were double-positive (Table S4). The *Chat* expressed CNs in STR-CP&ACB and PALm&PALv could be clearly discriminated according to the subtype feature genes (**Figure 6A**). The expression patterns of these feature genes were greatly coincident between the10X Visium and RCA-FISH results in STR-CP&ACB and PALm&PALv (Figure S8B; Figure S8C). We further compared the 10X and RCA-FISH results in terms of CNs’ sub-region distribution and feature gene expression. Both methods revealed comparable distribution ratios and similar expression patterns for CNs across different sub-regions (**Figure 6B**). For example, *Ndnf*, which is distinctively expressed in striatal CINs and was identified using STIF, also showed consistent results in RCA-FISH experiments (**Figure 6C**; **Figure 6D**). Particularly, we noticed *Parm1* was highly expressed in both Spot+ and Spot-in PALm&PALv, with STIF, it has been filtered out from the HEGs of BFCNs. Interestingly, in the RCA-FISH data, we found it was indeed highly expressed in PALm&PALv, but not co-expressed with BFCNs (Figure 6C; Figure 6D), which indicated the necessity of using STIF in 10X Visium data. In summary, the feature genes of the subtypes in CINs and BFCNs could be verified by RCA-FISH.

**Figure 6.**
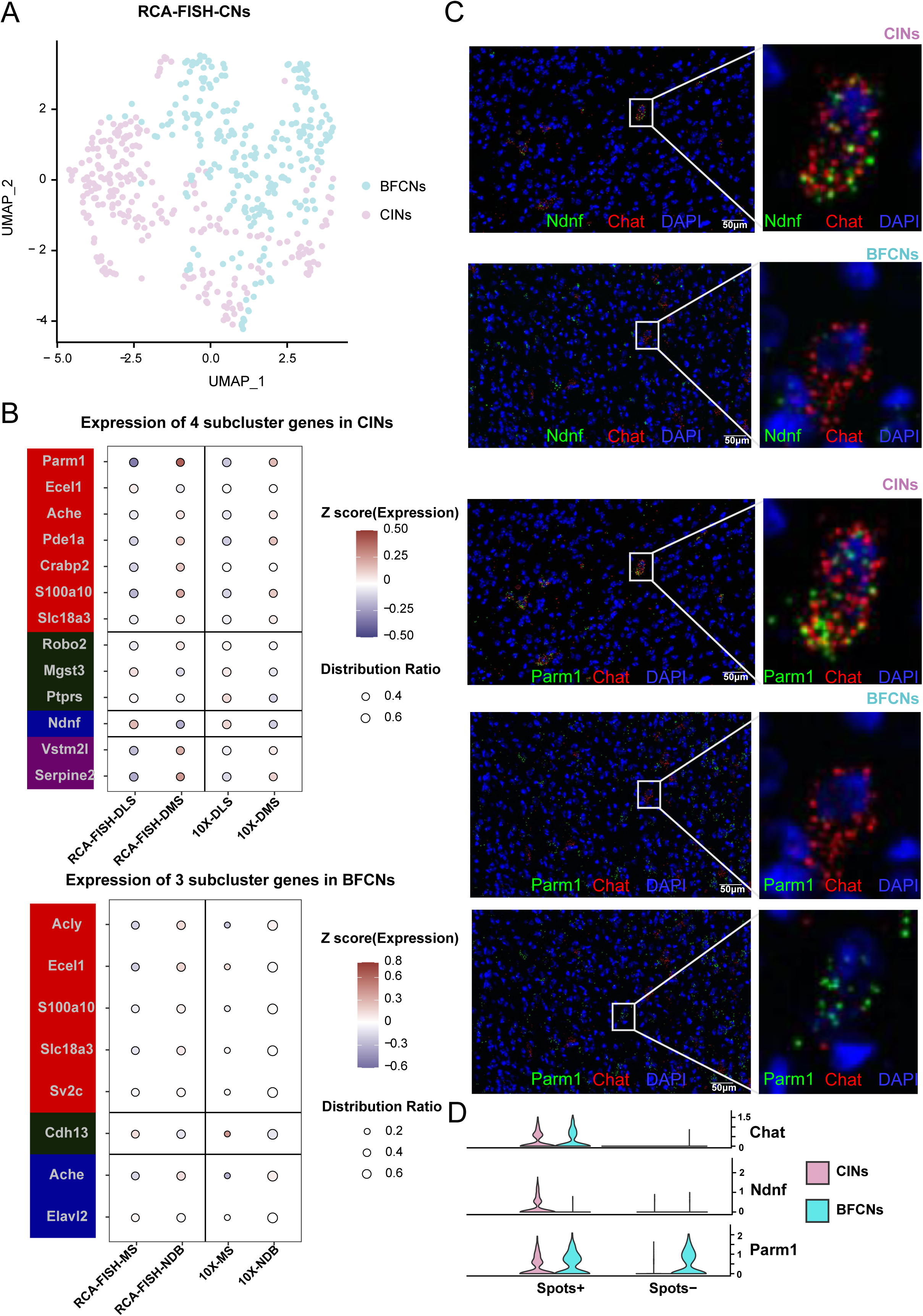
RCA-FISH of the subtype feature genes in striatal CINs and BFCNs. **A.** UMAP plot of the Chat-expressed CNs according to the subtype feature genes (CNs, n=487; CINs, n=240; BFCNs, n=247). **B.** The CNs’ sub-region distribution ratio and the subtype feature genes expression between the RCA-FISH and 10X Visium results in different brain regions (CNs, n=487). **C.** The RCA-FISH images of Ndnf and Parm1 in the CINs and BFCNs. **D.** The violin plot of Ndnf and Parm1 for "Spot+", "Spot-" in the CINs (n=408; 402) and BFCNs (n=295; 268).

## Discussion

The forebrain is responsible for a multitude of advanced functions, and its architectural subdivisions are considered functional modules that process various aspects (3). In brain regions, spatial gene expression is essential for understanding their identity and function. mRNAs and proteins are both important investigated objects in gene expression studies (72). Here, we performed MS–based proteome with laser capture microdissection and spatial transcriptome synchronously for comprehensive analysis of the forebrain regions.

Due to the current limitations of mass spectrometry detection, we have found that the expression of characteristic brain region proteins such as ChAT cannot be detected in tissue samples smaller than 936um × 20um (Figure S1 A). Under these constraints, we can only employ low-resolution sampling for spatial proteomics. Although this approach results in the loss of intra -TD information, it still allows for the effective exploration of characteristic genes in sub-anatomical brain regions like PALm&PALv, ACA and SS sub-region of Isocortex. With the further improvements of MS technology, we can expect resolutions to be improved, facilitating more accurate analysis of expression patterns. To ensure the homogeneity of protein data, we employed identical sampling sizes during MS detection.

The integration of transcriptomic and proteomic data led to the discovery of reliable molecular features in major brain regions and sub-regions of Isocortex. We identified 29 and 16 highly expressed genes with brain region specificity as molecular features. Of these, 18 are brain region feature genes and 16 are sub-region feature genes newly found in our study. These feature genes are closely linked to brain region functions. We also validated the newly discovered feature genes using Ortiz’s dataset, which revealed similar expression patterns in terms of expression and distribution.

Furthermore, we found that although the mRNA expression and protein abundance of HEGs and HEPs showed a low correlation, with Spearman coefficients ranging from 0.36 to 0.39, their participating functions exhibited high similarity based on GO analysis. To investigate the underlying factors, we visualized the profiles of enriched HEGs and HEPs on major synapse pathways in the Isocortex and STR-CP&ACB. We noticed that although they are enriched in the same functional pathway, the proportion of genes that exist in both mRNA and protein forms is low. Interestingly, we discovered that there is a difference in the ratio of enriched genes that exist in mRNA or protein forms between the Isocortex and STR-CP&ACB. In the Isocortex, the enriched genes are mainly in protein form in postsynaptic neurons, while in the STR-CP&ACB are mainly in mRNA form. This finding suggested that the neurons in the Isocortex were highly active, while the neurons in the STR-CP&ACB exhibited the capacity for rapid and precise motor control.

However, we observed that in the STR-CP&ACB and PALm&PALv, the spatial transcriptome and proteome data could not be clearly and simultaneously enriched into cholinergic synapses in the GO analysis. The issue might be attributed to the cell-blended environment and the relatively low proportion of CNs. The molecular characteristics of CNs could be obscured by other types of cells. To tackle this challenge, we introduced the STIF approach, which combined 10X Visium technology with IF signals. Notably, we observed that the CNs were predominantly distributed as single cells within each Visium spot. Leveraging the STIF approach, according to the expression specificity, we successfully extracted 12 feature genes of CINs and BFCNs from the cell-blended data. These feature genes are closely linked to the functions of CNs, 6 of which were newly discovered in our study. To validate these genes, we reanalyzed Munoz-Manchado’s scRNA-seq dataset and found that most of them were also specifically expressed in Chat-expressing cells. This work provided valuable insights into the characteristics of CNs and significantly improved the resolution of the analysis using 10X Visium data.

With the HEGs of CNs, we identified subtypes of CINs/BFCNs and their feature genes. These feature genes were validated with RCA-FISH. We observed that the subtypes of CINs exhibited distinct spatial distributions, showing a clear preference towards DLS or DMS. Interestingly, the distribution of BFCNs subtypes exhibited a mixed state among all the PALm&PALv regions. This is different from previous knowledge, which divided BFCNs subtypes into 4 groups according to their anatomical locations. However, our finding was consistent with the projection difference of BFCNs in MS/NDB (16), whose projection specificity is diverse between neighboring neurons. We also found that some subtype feature genes were related to neuronal projection. All of these findings suggest a potential association between neuronal gene expression and projection patterns.

In the future, by combining the spatial transcriptome data of CNs with the morphological features of sparsely labeled neuron projections through continuous slicing and imaging, a more comprehensive understanding of CNs from multiple perspectives could be achieved.

## Materials and methods

### Mice

C57BL/6 mice were obtained from Shanghai Slac Laboratory Animal Co., Ltd (SCXK (shanghai) 2017-0005). All experimental animal procedures and animal care were approved by the Animal Ethics Committee of Shanghai Institute of Nutrition and Health Institute (approval number: SINH-2023-ZGQ-1).

### Tissue collection for spatial transcriptomics

OCT-embedded whole brains were cryosectioned coronally to a thickness of 10µm (bregma: 0.74 to -0.48) using a freezing microtome (Leica 3050S). We layered tissue sections onto a spatially barcoded array to collect in situ 2D-RNAseq spatial transcriptomics (Lot #1000184 Visium Spatial Gene Expression Slide & Reagents Kit, 10X Genomics) and applied IF staining before tissue permeabilization. Each spatially barcoded array has 4992 spots, with a diameter of 55 µm and a center-to-center distance of 100 µm, over an area of 6.5 mm. One coronal section normally covers the area of 3000-3500 spots on the array. Each spot contains approximately 1 million barcoded reverse-transcription oligo (dT) primers and gets a global transcriptomic profile with a volume of 0.00002 mm3 (πr2h with r = 27.5 µm and h = 10 µm). After cryosection, all sections were stored at -80°C before proceeding with experiments.

### RNA integrity number

We collected 8 right and 8 left hemispheres from 2 different mice at 8 weeks of age. After taking 10 slices of 10-µm-thick cryosections, RNA was extracted and analyzed. RNA quality was checked by RNeasy Mini Kit (QIAGEN, Hilden, Germany) and Agilent 2100 Bio-analyzer with RNA nano chips (Agilent Technologies, Inc., Santa Clara, CA, USA). RIN values of the tissues were between 9.3 and 9.5.

### Spatial transcriptome sequencing

The spatial transcriptome sequencing was based on the Visium Technology Platform of 10x Genomics company (Lot #1000184 Visium Spatial Gene Expression Slide & Reagents Kit, 10X Genomics). The reagents and consumables in the experiment were provided by this platform, and the specific product numbers can be found at www.10xgenomics.com/products/spatial-gene-expression.

#### Optimization of the permeabilization time

Before using a new tissue for generating Visium Spatial Gene Expression libraries, the permeabilization time was optimized. Briefly, the Visium Spatial Tissue Optimization workflow included placing tissue sections on seven-capture areas on a Visium Tissue Optimization slide (from Visium Spatial Gene Expression Reagent Kit, 10x Genomics, PN-1000186). The sections were fixed, stained, and then permeabilized for different times. The mRNA released during permeabilization binds to oligonucleotides on the capture areas. Fluorescent cDNA was synthesized on the slide and imaged. The permeabilization time that results in maximum fluorescence signal with the lowest signal diffusion was optimal. If the signal was the same at two time points, then the longer permeabilization time was considered optimal. Once optimal conditions had been established, each cryosection was cut at 10 µm in thickness onto Visium Slide (from Visium Slide Kit) and processed immediately. In this study, the permeabilization time ranges from 3 to 18 min depending on the samples.

#### Tissue fixation, antibody staining, and imaging

Before tissue permeabilization, tissue sections on the Visium Slide (from Visium Slide Kit) were removed from −80°C and incubated 1min at 37°C, then the tissue sections were fixed in 40ml chilled methanol (MilliporeSigma) in a 50ml centrifuge tube by incubating 30 min at −20°C. For antibody staining, each tissue section was first incubated in 70µl Blocking buffer for 5min at room temperature, then incubated in 50µl 1:50 Alexa Fluor 488-anti-Chat Primary antibody solution for 30min at room temperature, followed by five washes with 100µl Wash buffer. (Blocking buffer, Primary antibody solution and Wash buffer were prepared according to 10X genomics protocol CG000312) RNase inhibitor and Ribonucleoside Vanadyl complex were used to prevent RNA degradation. Immerse the slide 20x in the 3X SSC buffer, add 200µl Mounting Medium (85% Glycerol, 2U/µl RNase Inhibitor) to cover the tissue sections on the slide uniformly. Then, the stained tissue sections were imaged with Zeiss Observer Z1 microscope immediately.

#### Tissue permeabilization and reverse transcription

For tissue permeabilization, the slides were first placed in the Slide Cassette (from the Visium Slide kit) for the optimal permeabilization time. A permeabilization enzyme (from the Visium Reagent kit) was used for permeabilizing the tissue sections on the slide for incubating for the predetermined permeabilization time. The polyadenylated mRNA released from the overlying cells was captured by the primers on the spots. After washing by 0.0.1×SSC (saline sodium citrate buffer, MilliporeSigma), RT Master Mix (provided in Visium Reagent kit) containing reverse transcription reagents was added to the permeabilized tissue sections in the Thermocycler Adaptor. Incubation with the reagents produces spatially barcoded full-length cDNA from polyadenylated mRNA on the slide.

#### Second strand synthesis and denaturation

After removing RT Master Mix (provided in Visium Reagent kit) from the wells, sections were incubated in 0.08 M KOH for 5 min and washed by Buffer EB (QIAGEN). Then, the Second Strand Mix (provided in Visium Reagent kit) was added to the tissue sections. On the slide to initiate second strand synthesis on the Thermocycler Adaptor. This is followed by denaturation and transfer of the cDNA from each capture area to a corresponding tube for amplification and library construction. The slides were washed by Buffer EB and incubated in 0.08 M KOH for 5 min. Then, samples from each well were transferred to a corresponding tube containing tris-HCl (1 M; pH 7.0) in eight-tube strip for amplification and library construction.

#### cDNA amplification and quality control

Denaturation sample of 1 µl was transferred to the quantitative polymerase chain reaction (qPCR) plate well containing the qPCR Mix [nuclease-free water + KAPA SYBR FAST qPCR Master Mix (KAPA Biosystems) + cDNA Primers (from Visium Reagent kit)]. The Cq value for each sample was recorded after qPCR. For cDNA amplification, cDNA Amplification Mix (from Visium Reagent kit) was added to the remaining sample from denaturation. Then, the product was incubated in Thermocycler Adaptor for a cycle. For cDNA Cleanup–SPRIselect, 60 µl of SPRIselect reagent (Beckman Coulter) was added to each sample and incubated for 5 min at room temperature. The sample was repeatedly adsorbed by the magnet·High, washed with ethanol (MilliporeSigma) and Buffer EB and transferred to a new tube strip. Then, we ran 1 µl of sample on an Agilent Bioanalyzer High Sensitivity chip (Agilent, catalog no. 50674626) for cDNA quality control (QC) and quantification.

#### Visium spatial gene expression library construction

Enzymatic fragmentation and size selection were used to optimize the cDNA amplicon size. P5, P7, i7, and i5 sample indexes and TruSeq Read 2 (read 2 primer sequence) were added via End Repair, A-tailing, Adaptor Ligation, and PCR. The final libraries contain the P5 and P7 primers used in Illumina amplification. Library construction was performed with Library Construction Kit (10x Genomics, catalog no. PN-1000190).

#### Fragmentation, End Repair, and A-tailing

Only 10 µl of purified cDNA sample from cDNA Cleanup was transferred to a tube strip. Buffer BE and Fragmentation Mix (from Library Construction kit) were added to each sample, and fragmentation was performed in a thermal cycler. After fragmentation, 30 µl of SPRIselect reagent (0.6×) was added to each sample and incubated for 5 min at room temperature. The sample was repeatedly adsorbed by the magnet·High, washed with ethanol and Buffer EB, and transferred to a new tube strip.

#### Adaptor ligation

Adaptor Ligation Mix (from Library Construction kit; 50 µl) was added to each 50 µl of sample and incubated in a thermal cycler.

#### Postligation cleanup–SPRIselect

SPRIselect reagent (0.8×) was added to each sample and incubated for 5 min at room temperature. The sample was repeatedly adsorbed by the magnet·High, washed with ethanol and Buffer EB, and transferred to a new tube strip.

#### Sample index PCR

Amp Mix (from Library Construction kit; 50 µl) and an individual Dual Index TT Set A (10x Genomics, catalog no. PN-1000215, 20 µl) were added to each 30 µl of sample and incubated in a thermal cycler.

#### Post–sample index PCR double-sided size selection–SPRIselect

SPRIselect reagent (0.6×) was added to each sample and incubated for 5 min at room temperature. After adsorbing by the magnet·High, 150 µl of supernatant was transferred to a new tube strip. Then, 20 µl of SPRIselect reagent (0.8×) was added to each sample and incubated for 5 min at room temperature. After adsorbing by the magnet·High and supernatant removed, samples were washed with ethanol and Buffer EB and transferred to a new tube strip. We ran 1 µl of the sample (1:10 dilution) on an Agilent Bioanalyzer High Sensitivity chip.

#### Sequencing

A Visium Spatial Gene Expression library comprises standard Illumina paired-end constructs that begin and end with P5 and P7. The 16–base pair (bp) spatial barcode and 12-bp UMI are encoded in Read 1, while Read 2 is used to sequence the cDNA fragment. i7 and i5 sample index sequences are incorporated. TruSeq Read 1 and TruSeq Read 2 are standard Illumina sequencing primer sites used in paired-end sequencing.

### Laser capture microdissection of the brain section

Laser capture microdissection was performed on 8 extra hemispheres immediately adjacent to the 8 brain sections used for Visium Spatial RNA-seq individually (with ABI Arcturus XT). Tissue domains with an average diameter 936µm were dissected using ABI Arcturus XT. On average, 21 tissue domains (TDs) were dissected from one hemisphere. After microdissection, all tissue domains were stored at -80°C individually before proceeding with trace mass spectrometry. In the meantime, the dissection positions were recorded by imaging. To test the minimum reasonable thickness of the TD, we used ChAT as a marker for the TD from MS-NDB. Only when the ChAT could be detected by HPLC-MS/MS, the thickness considered to be valid. 20µm was the minimum thickness that could be identified by ChAT (Figure S8A).

### Liquid chromatography tandem mass spectrometry analysis of peptide mixture

Samples were analyzed on Q Exactive HF mass spectrometer (Thermo Fisher Scientific, Rockford, IL, USA) coupled with an Easy-nLC 1200 nanoflow LC system (Thermo Fisher Scientific). In brief, 0.5 μg of peptide mixture resolved in buffer A (0.1% formic acid (FA)) were loaded onto a 2-cm self-packed trap column (150 μm inner diameter, C18, 3 μm) using buffer A and separated on a 100-μm-inner-diameter column with a length of 12 cm (C18, 1.9 μm) over a 75-min gradient (buffer A, 0.1% FA in water; buffer B, 0.1% FA in ACN) at a flow rate of 600 nl/min (0–10 min, 4–15% B; 10–60 min, 15–30% B; 60–69 min, 30–50% B; 69–70 min, 50–100% B; and 70–75 min,100% B). The Orbitrap Fusion was set to the OT–IT mode. For a full mass spectrometry survey scan, the target value was 5 × 105 and the scan ranged from 300 to 1,400 m/z at a resolution of 120,000 and a maximum injection time of 50 ms. For the MS2 scan, a duty cycle of 3 s was set with the top-speed mode. Only spectra with a charge state of 2–6 were selected for fragmentation by higher-energy collision dissociation with a normalized collision energy of 32%. The MS2 spectra were acquired in the ion trap in rapid mode with an AGC target of 1lJ×lJ104 and a maximum injection time of 35 ms.

### RCA-FISH for brain sections

The RNA RCA-FISH was designed and improved from the STARmap protocol (73). And the experimental protocol was modified based on STARmap. Briefly, the brain slices were fixed with 4% PFA at RT for 5 min, and transferred them in permeabilization buffer comprising 2×SSC, 2% v/v sodium dodecyl sulfate (SDS), 0.5% Triton-X-100 for 2min, then stored in ethanol at 4℃ overnight. The next day, the brain slices incubated in 1× hybridization buffer (2× SSC, 35% formamide, 1% Triton-X-100, 0.2% SDS, 20mM RVC, 0.1 mg/ml tRNA and pooled SNAIL probes at 50 nM per oligo) at 37℃ overnight. The brain slices were ligated by T4 DNA Ligation mixture (0.2 U/ul T4 DNA Ligase supplemented with 0.2U/ul RRI) and amplified by Phi29 DNA polymerase mixture (0.2 U/ul Phi29 DNA polymerase, 250 uM dNTP, 0.2 U/ul RRI and 10nM 5-(3-aminoally1)-dUTP), then modified by Acrylic acid NHS ester and embedded with Acrylamide hydrogel. For simultaneous staining IF with RCA-FISH for Chat, the tissue gel was incubated in 1:50 anti-ChAT overnight and imaged with Olympus VS200. After IF imaging, the tissue gel was digested with Proteinase K to remove the IF and to reduce the background signal, simultaneously. For RNA read-out detection, we used multiple round imaging procedures with Olympus VS200 or KF-FL-005, Ningbo KFBIO. For each round read-out probe hybrid, 3 probes were hybrid with 2×SSC, 10% v/v ethylene carbonate incubated at RT for 30 min. After imaging, samples were incubated with buffer with 10% TCEP (Tris (2-carboxyethyl) phosphine hydrochloride) and washed with 2×SSC to remove the fluorescent signal that coupled with disulfide bond in the fluorescent oligos. The multiple round images were registered and analyzed with Arivis.

### Quantification and Statistical Analysis

#### Multi-omics data registration

For a group of adjacent brain sections, the first section is a brain section on Visium chip with anti-ChAT immunofluorescence labeling used for spatial transcriptomics, while the immediately adjacent section is a laser microdissected brain section used for spatial proteomics. Both the spots of spatial transcriptomics and the TDs of spatial proteomics contain positional information. The precise locations of the TDs were documented during microdissection by referencing the cutting marks using bright-field imaging. To associate anti-ChAT signal, spatial transcriptome and proteome data and analyze them synchronously, the brain section on Visium chip with anti-ChAT IF was first registered with the immediately adjacent microdissected brain section according to the outline of the brain sections. The microdissected –Visium chip registered image was then imported to the Loupe Browser according to the fiducial frames, which positioned the Visium spots. This process allowed the TDs on the microdissected brain section to be accurately associated with the Visium spots and the anti-ChAT signal on the Visium chip.

#### Brain areas annotation

Standard 2D mouse brain reference atlases, e.g., the Allen Reference Atlas (ARA) were used to annotate the brain areas (74). The current standard 2D atlas annotations were drawn on the IF brain section images. The anti-CHAT-488 stained IF signal, the tissue structure shown by spontaneous fluorescence and the Visium Spatial RNA-seq cluster output were used to fine - tune the boundary of the brain areas. SMDB were used for 3D visualization of CN subclusters (75).

#### Raw sequencing data processing

The Visium Spatial RNA-seq output and the fluorescence microscope images were analyzed by Space Ranger (version 1.1.0) with the GRCh38 v86 genome assembly as reference to detect tissue, align reads and generate gene-spot matrices. This pipeline combined Visium-specific algorithms with the widely used RNA-seq aligner STAR. Seurat package (version 3.1.3) was used for gene-spot matrices processing and downstream analysis in R besides custom scripts (76). Quality control was performed for each gene-spot matrix of brain slices, the spots with detected gene numbers under 200 were removed while the genes with fewer than 10 read counts or expressed in fewer than 2 spots were removed. Normalization across spots was performed with the "SCTransform" function. Highly variable genes (HVG) were selected using the "FindVariableFeatures" function in Seurat with the ‘vst’ setting for 5,000 features; Feature information was then scaled, and dimensionality reduction and clustering was performed with PCA at resolution 0.8 with the first 30 PCs. Spatial feature expression plots were generated with the "SpatialFeaturePlot" function.

#### Quality control of the mass spectrometry platform

For the quality control of the performance of mass spectrometry, the HEK293T cell (National Infrastructure Cell Line Resource) lysate was measured as the quality-control standard. The quality-control standard was digested and analyzed using the same method and conditions as the mouse samples. A Pearson’s correlation coefficient was calculated for all quality-control runs in the statistical analysis environment R (version 3.6.1). The correlations of the protein quantification results were kept for repeated samples with 0.88∼0.94 between HEK293T cells (Figure S8B). The result demonstrated the consistent stability of the mass spectrometry platform.

#### Protein identification by MaxQuant-based database searching

The tandem mass spectra were searched against the human UniProt database (version 20210225, 17,193 sequences) using MaxQuant (version 1.6.7) (77). Trypsin was selected as the proteolytic enzyme and two missed cleavages sites were allowed. Cysteine carbamidomethylation was set as the fixed modification. The oxidation of M and acetylation of the protein N-terminal were set as the variable modifications. The false discovery rates of the peptide-spectrum matches (PSMs) and proteins were set to less than 1%. Protein identification required at least one unique or razor peptide per protein group. Quantification in MaxQuant was performed using the label free quantification (LFQ) algorithm. Briefly, MaxLFQ is an intensity determination and normalization procedure that is fully compatible with any peptide or protein separation before LC-MS analysis. LFQ of 165 samples were extracted from the MaxQuant result files to represent the final expression of a particular protein across samples, resulting in a 3,978 × 165 protein-expression matrix. Then, the expression matrix was log2-transformed and used in all quantitative analysis.

#### Principal component analysis in all forebrain regions

For brain region analysis, a merged genes-spots matrix of eight coronal slices need to be generated with the “merge” Seurat function. The merged matrix was normalized with "SCTransform" function prior to PCA and UMAP on the first 30 PCs. We extracted expression data for genes and proteins to conduct principal component analysis with "prcomp" in R, respectively. Using the top two principal components, we looked at the relevance of expression across brain regions in terms of transcriptome and proteome.

#### Identification of highly expressed genes and proteins from forebrain regions

Differential gene expression analysis was performed using the "FindMarkers" function in Seurat (version 3.1.3). The R Limma package (version 3.56.1) was used to process proteomic data (78). The differences between each brain region and the other brain regions (1: 4 other regions in major brain region and 1:2 in sub-regions) were assessed using multivariate linear models. We selected genes and proteins with log2FC > 0.25 and false discovery rate (FDR) < 0.05 to screen HEGs and HEPs of 5 major brain regions and sub-regions in Isocortex.

#### Functional enrichment analysis of forebrain regions

The package of clusterProfiler (version 4.8.1) was used to evaluate the biological pathways and processes of HEGs and HEPs related to brain regions (79), we performed gene ontology (GO) enrichment analysis examining biological components, molecular functions, biological processes, and KEGG pathways in major brain regions (80, 81). HEGs and HEPs were labeled by pathview (version 1.2.3) to show their up and downstream positions in five main synapse pathways.

#### Identifying gene sets with high correlation within mRNA expression and protein abundance in brain regions

The package of randomForest (version 4.7) was used to identify the gene sets. The correlation of sample pairs was categorized into two classes: "high" and "low", based on which a classification model was constructed. Using the Random Forest algorithm, we calculated feature importance scores for each gene to evaluate its contribution to the model’s predictive power. Subsequently, genes were ranked according to these feature importance scores, highlighting those genes that have the greatest impact on the observed correlations.

#### Spatial-transcriptome and IF joint analysis (STIF) for cholinergic neuron molecular characteristic profile

The mRNA profiles in each 10X Visium spot are admixture of multiple cells and the CNs are 1/7 of the cells in the Visium spot, so it is challenging to extract the molecular features of the CNs. We integrated spatial-transcriptome data with IF signals and developed a strategy with three main processes called Spatial-transcriptome and IF joint analysis (STIF). **Process I: Label spots as "Spot+" and "Spot-".** The 10X Visium spots containing anti-ChAT IF signals were selected to be "Spot+", while the spots immediately adjacent to the "Spot+", in and around which did not contain any anti-ChAT IF signal to be the "Spot-". The spots belonging to neither "Spot+" nor "Spot-" were labeled as "Others" to indicate the background. **Process II: Differential expression analysis of "Spot+" and "Spot-".** We compared the transcriptomics profiles between "Spot+" and "Spot-" in the STR-CP&ACB and the PALm&PALv respectively (Figure 4B). Differential gene expression analysis was performed using the "FindMarkers" function in Seurat. A wilcoxon rank-sum statistic test between "Spot+" and "Spot-" in STR-CP&ACB and PALm&PALv was performed respectively to obtain the molecular characteristics of CINs and BFCNs. HEGs with both Bonferroni adjusted P value <0.05 and log2FC > 0.25 were considered to be statistically significant and potentially to be cholinergic neuron features. **Process III: Subtyping of CINs and BFCNs in STR-CP&ACB and PALm&PALv.** Well-separated "Spot+" in STR-CP&ACB (n=274) or PALm&PALv (n=161) were clustered with CINs or BFCNs HEGs respectively to identify cholinergic neuron subtypes, with louvain algorithm using "FindClusters" function in Seurat. The HEGs identified in various subclusters were considered as subtype-specific features of CINs and BFCNs. **Another comparing strategy.** The HEGs of the "Spot+" between the STR-CP&ACB and the PALm&PALv and the HEGs of the "Spot-" between the STR-CP&ACB and the PALm&PALv were firstly identified (Figure S6B). Afterwards, the HEGs of CINs and BFCNs were identified by comparing the "Spot+ HEGs" and the "Spot-HEGs".

#### Clustering result evaluation of Spot+, Spot-in STR-CP&ACB and PALm&PALv with different gene sets

ARI (Adjusted Rand Index) scores were calculated through "metrics.adjusted_rand_score" function with Scikit-learn module of Python3 to evaluate clustering result of Spot+ and Spot-in STR-CP&ACB and PALm&PALv with different gene sets respectively.

#### Normalization of the distribution proportions of CNs across subclusters

The distribution proportions of CNs across subclusters were normalized in a two-step process to enable direct comparison across subclusters and brain subregions. Firstly, the proportion of each subcluster within each brain region was determined based on its spot count relative to the total spot count in that region. Secondly, these proportions were standardized by expressing the proportion of each subcluster in a specific region as a fraction of its total proportion across all regions, yielding normalized proportions.

## Authors’ contributions

Y.C., Q.L. and G.Z. conceived of the project, supervised the study and revised the manuscript. Y.C. and N.C. performed the spatial transcriptomic and proteomic experiments. M.H. and J.M. analyzed the spatial transcriptomic and proteomic data and wrote the manuscript. Y.C., J.Y., D.C., M.H. and N.C. performed the RNA RCA-FISH experiments, imaging, registration and analysis.

## Competing interests

All other authors declare they have no competing interests.

## Acknowledgements

Funding for the study was supported by grants from the National Major Scientific Instrument and Equipment Development Project of NSFC (81827901), the Strategic Priority Research Program of the Chinese Academy of Sciences (XDB38030100). The work was also supported by Shanghai Municipal Science and Technology Major Project 2023SHZDZX02 and GTP project.

## Data Availability

All data can be viewed in NODE (http://www.biosino.org/node) by pasting the accession OEP004212 into the text search box or through the URL: http://www.biosino.org/node/project/detail/OEP004212.

## Supplementary material

**Figure S1 Quality control for mass spectrometry experiment.**

**A.** Images of the tissue domains and brain sections after laser capture microdissection. **B.** Quality control of mass spectrometry using tryptic digest of HEK293T cells. The numbers represent the pairwise Spearman’s correlation coefficients of the samples (cells, n=9).

**Figure S2 Analysis and validation of the in situ mRNA and protein expression in the brain regions.**

**A.** Uncombined and combined brain regions. **B.** Heat map of HEGs and HEPs across the 5 brain regions (log2-fold-change (Log2FC)> 0.25 and FDR<0.05). **C.** Scatter plot of log2FC versus log2 transcript per million (TPM) of the top 20 HEGs (left) and log2FC versus log2 LFQ of the top 20 HEPs (right) in the 5 brain regions. **D.** The mRNA spatial distribution of 29 region-specific feature genes (left). The boxplot of the mRNA expression and protein abundance of these 29 genes in relevant brain regions (Isocoretx, TDs=48; OLF-PIR, TDs=4; STR-CP&ACB, TDs=30; PALm&PALv, TDs=3; Hypothalamus, TDs=4; wilcoxon test, P<0.05) (right). **E.** The scatter plot of Log2FC for 29 feature genes in Ortiz dataset and our data. **F.** The mRNA spatial distribution of 29 region-specific feature genes in Ortiz’s dataset.

**Figure S3 mRNA spatial distribution and protein expression of 16 subregion-specific genes in the Isocortex.**

**A.** The mRNA spatial distribution of 16 subregion-specific genes in ACA and SS (left). The boxplot of the mRNA expression and protein abundance of these 16 genes in the Isocortex (ACA, TDs=7; SS, TDs=21; wilcoxon test, P<0.05) (right). **B.** The scatter plot of Log2FC for 16 sub-region feature genes in Ortiz dataset and our data for validation. **C.** The mRNA spatial distribution of ACA and SS region-specific genes in Ortiz’s dataset.

**Figure S4 Analyze the relationship of the mRNA expression and protein abundance in the multi-omics data.**

**A.** Correlation plot of the mRNA expression (left) or protein abundance (right) across the 5 brain regions respectively. **B.** Fisher tests of GO enrichment between HEGs and HEPs across 5 brain regions.

**Figure S5 The KEGG pathviews of mRNAs and proteins in the synapse pathways in the Isocortex and STR-CP&ACB.** Glutamatergic and GABA-ergic synapse in the Isocortex; the dopaminergic, cholinergic and GABA-ergic synapse in the STR-CP&ACB. The proportion of different expression forms in the presynapse site. The genes highly expressed mRNAs were colored by purple, highly expressed proteins were colored by green and highly expressed mRNAs and proteins synchronously were colored by yellow.

**Figure S6 Superiority of STIF, expression and spatial distribution of CNs feature genes.**

**A.** Bar plot of cell numbers counted in single spots on 8 coronal sections of the mouse brain (10 spots per section were counted, total number=40). **B.** The diagram of another comparing strategy:The HEGs of the "Spot+" between the STR-CP&ACB and the PALm&PALv (n=408;295), and the HEGs of the "Spot-" between the STR-CP&ACB and the PALm&PALv (n=402;268) were first identified. The HEGs of CINs and BFCNs were identified by comparing the "Spot+ HEGs" (n=961) and the "Spot-HEGs" (1090), with wilcox test, FDR<0.05, log2FC>0.25. **C.** UMAP plots of the "Spot+" and "Spot-" in STR-CP&ACB (n=408; 402) and PALm&PALv (n=295; 268) with 130 HEGs from another comparing strategy and top 2000 highly variable genes from Seurat workflow. **D.** The boxplot of ARI scores, which were derived from clustering results corresponding to 77, 2000 and 130 genes (FDR=0.05; 0.05; 0.0019 respectively). **E.** Dot plot of HEGs in the striatal CINs or BFCNs with their log2FC value, log2FC>0.25. 12 genes with distribution specificity were labeled. **F.** The spatial distribution of CNs feature genes on the brain sections. **G.** The violin plots of 9 specific expressed genes of Chat+ neurons (n=60) in Munoz-Manchado’s scRNA-seq dataset.

**Figure S7 2D spatial visualization of the striatal CINs/BFCNs subclusters and their feature genes expression.**

**A.** The mRNA spatial distribution of Cnr1 and Crym on the brain sections. **B.** The 2D spatial visualization of STR CINs subclusters in CP and ACB across 8 brain coronal sections (IF1-8), the "Spot+" of different STR CINs subclusters were indicated by red, green, blue and purple. **C.** The 2D spatial visualization of BFCNs subclusters in MS, MA, SI and NDB across 8 brain coronal sections (IF1-8), the "Spot+" of different BFCNs subclusters were indicated by red, green and blue. **D.** UMAP plots of 13 feature genes in STR CINs subclusters, these subcluster feature genes were indicated by red, green, blue and purple respectively. **E.** UMAP plots of 8 feature genes in BFCNs subclusters, these subcluster feature genes were indicated by red, green and blue respectively.

**Figure S8 The 10X expression and RCA-FISH images of the striatal CINs and BFCNs subtype feature genes.**

**A.** Anti-ChAT and Chat-RCA-FISH images of CINs and BFCNs. **B.** The violin plot of the subtype feature genes for "Spot+", "Spot-" in the CINs (n=408; 402) and BFCNs(n=295; 268). **C.** The RCA-FISH images of the striatal CINs and BFCNs subtype feature genes.

**Table S1** HEGs, HEPs and molecular feature genes in the 5 brain regions and sub-regions of Isocortex

**Table S2** GO and KEGG enrichment results in the 5 brain region; glutamatergic, GABA-ergic, dopaminergic and cholinergic synapse pathways related genes in the Isocortex and STR-CP&ACB

**Table S3** The HEGs list and molecular feature genes of striatal CINs, BFCNs and their subclusters.

## Notes

### Competing Interest Statement

The authors have declared no competing interest.

