## Supplementary material for "Mouse Forebrain Region-specific Molecular Characteristics and Cholinergic Neurons Subtyping: An Integrated Analysis Based on Spatial Multi-omics": Figure S1-S8,Table S1-S4: Figure S1.pdf

A

### Test the minimum reasonable thickness of the TD

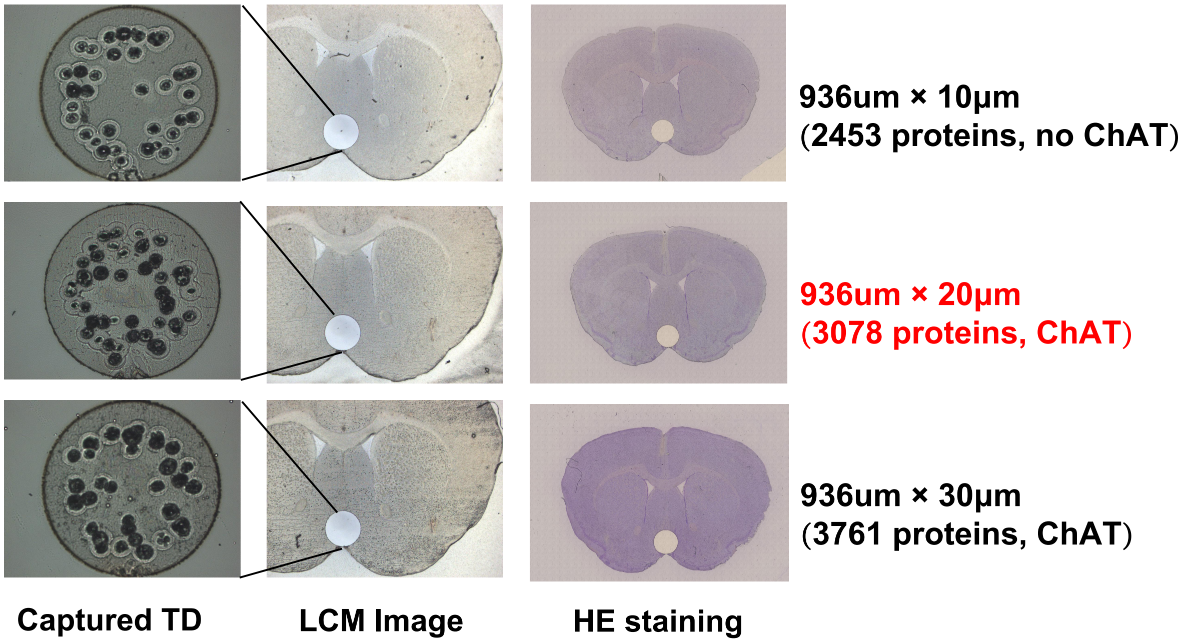


B

### Spearman's correlation coefficients of HEK293T controls experiments

Range of correlation: 0.88-0.94

| | | | | | | | | |
|---|---|---|---|---|---|---|---|---|
| Sample 1 | 0.93 | 0.91 | 0.90 | 0.91 | 0.91 | 0.89 | 0.89 | 0.90 |
| 0.93 | Sample 2 | 0.90 | 0.90 | 0.90 | 0.90 | 0.88 | 0.89 | 0.90 |
| 0.91 | 0.90 | Sample 6 | 0.94 | 0.93 | 0.93 | 0.90 | 0.89 | 0.90 |
| 0.90 | 0.90 | 0.94 | Sample 7 | 0.94 | 0.93 | 0.89 | 0.90 | 0.90 |
| 0.91 | 0.90 | 0.93 | 0.94 | Sample 4 | 0.94 | 0.91 | 0.90 | 0.91 |
| 0.91 | 0.90 | 0.93 | 0.93 | 0.94 | Sample 5 | 0.89 | 0.89 | 0.90 |
| 0.89 | 0.88 | 0.90 | 0.89 | 0.91 | 0.89 | Sample 3 | 0.89 | 0.89 |
| 0.89 | 0.89 | 0.89 | 0.90 | 0.90 | 0.89 | 0.89 | Sample 8 | 0.90 |
| 0.90 | 0.90 | 0.90 | 0.90 | 0.91 | 0.90 | 0.89 | 0.90 | Sample 9 |
