## Supplementary material for "Mouse Forebrain Region-specific Molecular Characteristics and Cholinergic Neurons Subtyping: An Integrated Analysis Based on Spatial Multi-omics": Figure S1-S8,Table S1-S4: Figure S4.pdf

A

### Correlation coefficients of the five brain regions in the transcriptome and proteome

| | Protein-Isocortex | Protein-OLF-PIR | Protein-STR-CP&ACB | Protein-PALm&PALv | Protein-Hypothalamus |
|---|---|---|---|---|---|
| Protein-Isocortex | 0.92 | 0.89 | 0.85 | 0.83 | 0.84 |
| Protein-OLF-PIR | | 0.93 | 0.86 | 0.84 | 0.84 |
| Protein-STR-CP&ACB | | | 0.91 | 0.81 | 0.80 |
| Protein-PALm&PALv | | | | 0.92 | 0.86 |
| Protein-Hypothalamus | | | | | 0.89 |

| | mRNA-Isocortex | mRNA-OLF-PIR | mRNA-STR-CP&ACB | mRNA-PALm&PALv | mRNA-Hypothalamus |
|---|---|---|---|---|---|
| mRNA-Isocortex | 0.64 | | | | |
| mRNA-OLF-PIR | 0.60 | 0.62 | | | |
| mRNA-STR-CP&ACB | 0.51 | 0.52 | 0.66 | | |
| mRNA-PALm&PALv | 0.54 | 0.56 | 0.50 | 0.64 | |
| mRNA-Hypothalamus | 0.50 | 0.52 | 0.45 | 0.63 | 0.65 |

B

### Fisher test of GO enrichment between the transcriptome and proteome

| | Protein-Isocortex | Protein-OLF-PIR | Protein-STR-CP&ACB | Protein-PALm&PALv | Protein-Hypothalamus |
|---|---|---|---|---|---|
| mRNA-Isocortex | 1039 (1.43E-130) | 275 (7.91E-19) | 690 (5.14E-26) | 165 (0.17) | 249 (1.77E-3) |
| mRNA-OLF-PIR | 459 (5.95E-25) | 204 (3.86E-40) | 336 (4.86E-11) | 69 (0.17) | 78 (1.77E-3) |
| mRNA-STR-CP&ACB | 913 (1.94E-29) | 229 (0.05) | 859 (6.69E-104) | 158 (0.17) | 224 (1.77E-3) |
| mRNA-PALm&PALv | 352 (3.08E-08) | 85 (0.57) | 302 (9.67E-13) | 202 (1.97E-74) | 130 (7.40E-07) |
| mRNA-Hypothalamus | 247 (0.81) | 65 (0.67) | 208 (0.15) | 93 (7.49E-08) | 190 (7.18E-44) |
