## Supplementary material for "Mouse Forebrain Region-specific Molecular Characteristics and Cholinergic Neurons Subtyping: An Integrated Analysis Based on Spatial Multi-omics": Figure S1-S8,Table S1-S4: Figure S5.pdf

### Glutamatergic synapse in Isocortex

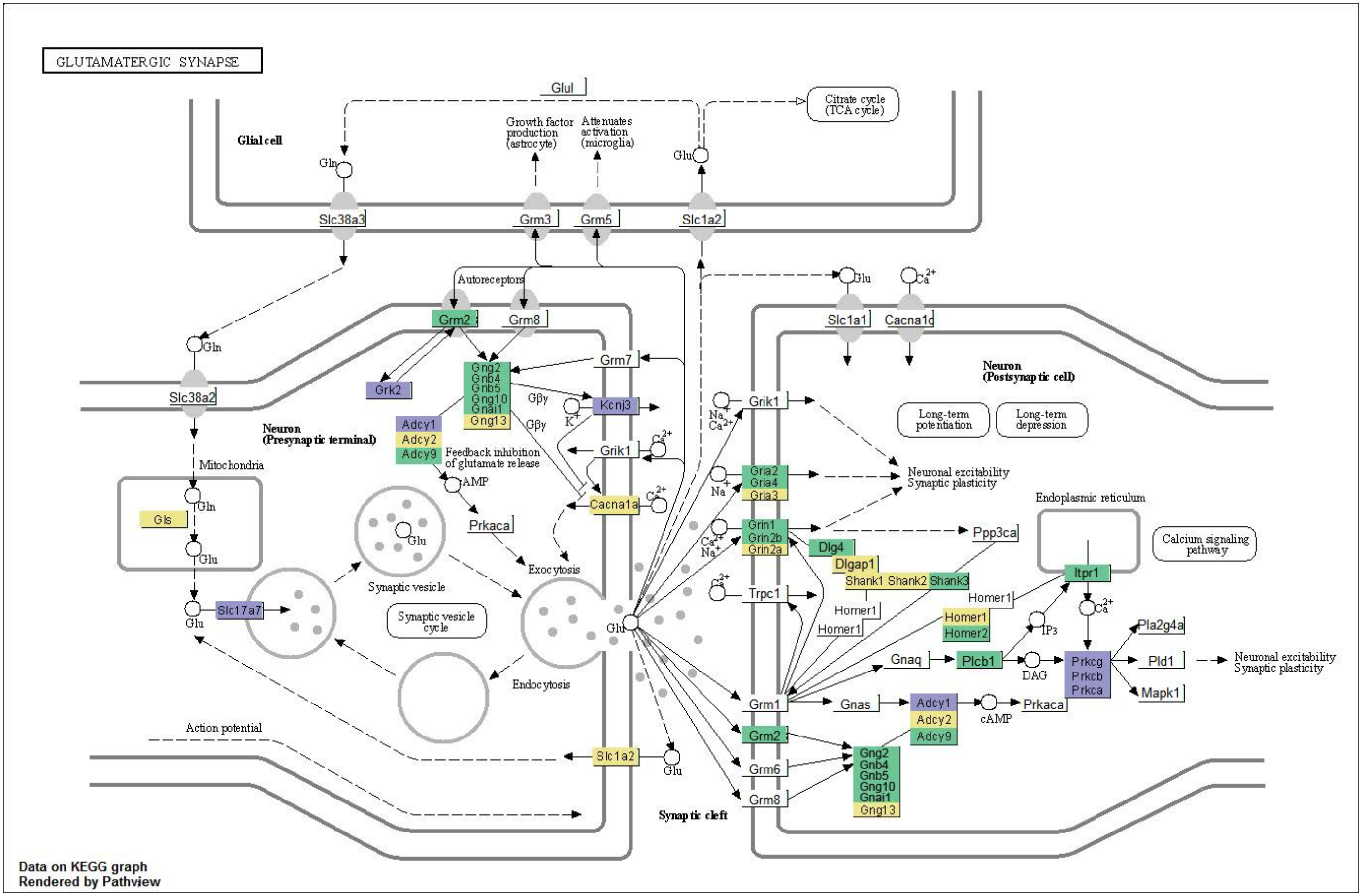


**Pre:** Slc17a7 Grk2 Adcy1 Kcnj3
**Pre:** Gls Adcy2 Gng13 Cacna1a Slc1a2
**Pre:** Adcy9 Grm2 Grm3 Gng2 Gnb4 Gnb5 Gng10 Gnai1

**Post:** Prkcg Prkcb Prkca
**Post:** Gria3 Grin2a Dlgap1 Shank1 Shank2 Homer1 Gng13 Adcy2
**Post:** Gria2 Gria4 Grin1 Grin2b Dlg4 Shank3 Homer2 Plcb1 Itpr1 Adcy9 Gng2 Gnb4 Gnb5 Gng10 Gnai1

### GABAergic synapse in Isocortex

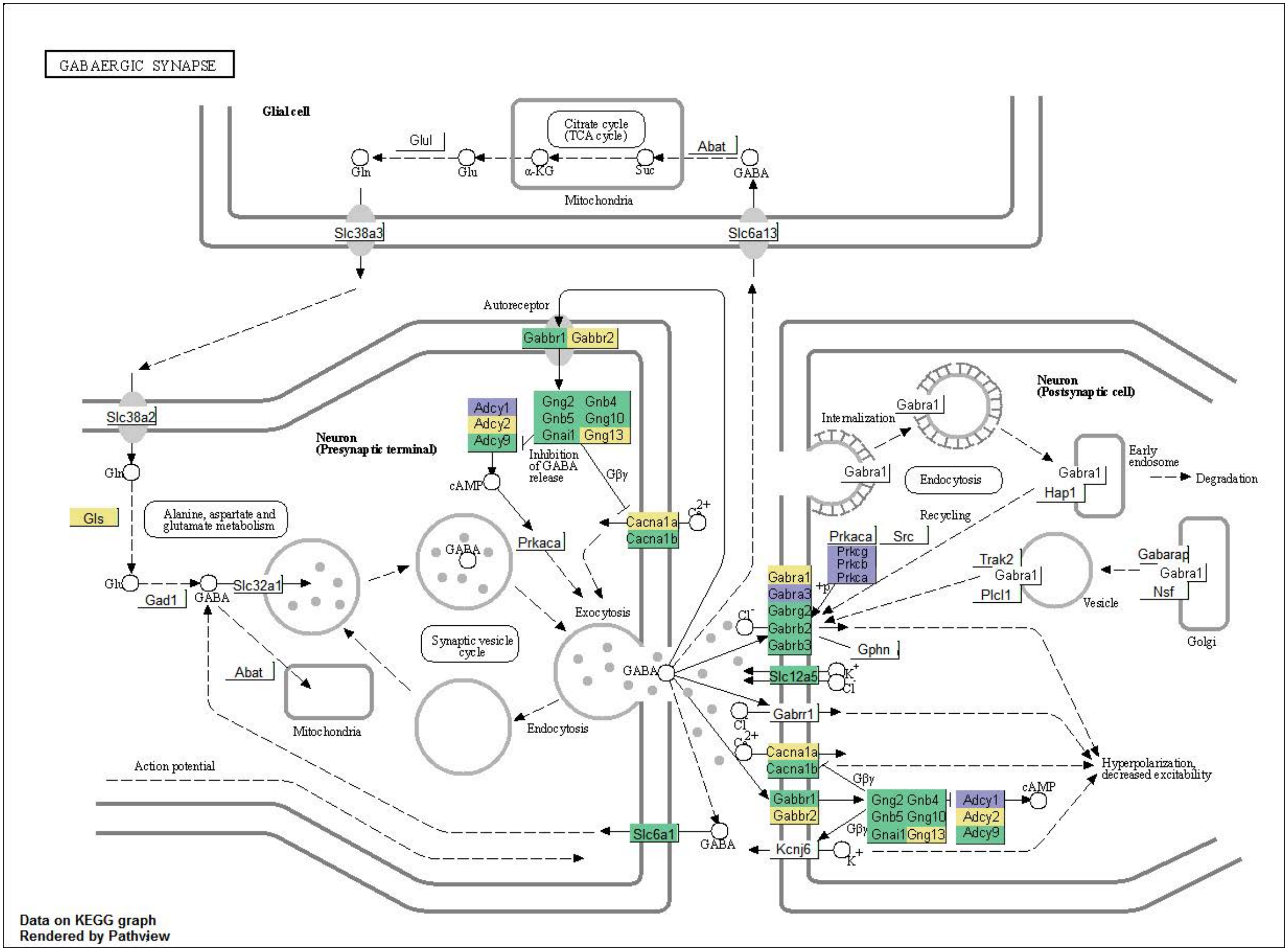


**Pre:** Adcy1
**Pre:** Gls Adcy2 Cacna1a Gng13 Gabra1 Gabbr2
**Pre:** Gng2 Gnb4 Gnb5 Gng10 Gnai1 Adcy9 Cacna1b Slc6a1

**Post:** Prkcg Prkcb Prkca Gabra3
**Post:** Adcy2 Cacna1a Gng13 Gabra1 Gabbr2
**Post:** Gng2 Gnb4 Gnb5 Gng10 Gnai1 Adcy9 Cacna1b Slc12a5 Gabrg2 Gabbr1 Gabrb2 Gabrb3

### Dopaminergic synapse in STR-CP&ACB

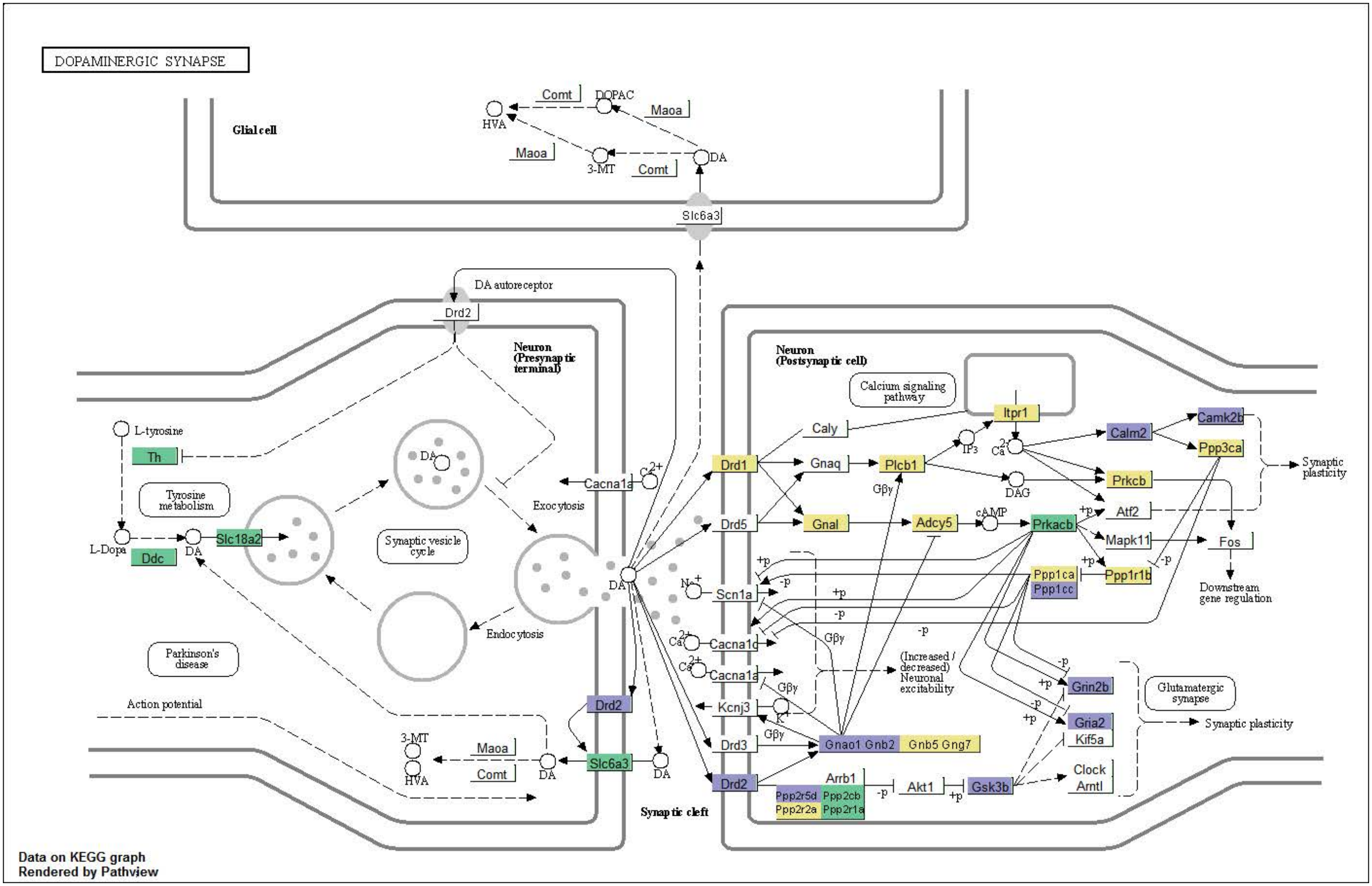


**Pre:** Drd2

**Pre:**

**Pre:** Th Ddc Slc18a2 Slc6a3

**Post:** Drd2 Gnao1 Gnb2 Ppp2r5d Gsk3b Grin2b Gria2 Ppp1cc Calm2 Camk2b

**Post:** Drd1 Gnal Plcb1 Adcy5 Itpr1 Prkcb Ppp1r1b Ppp3ca Gnb5 Gng7 Ppp1ca Ppp2r2a

**Post:** Ppp2cb Ppp2r1a Prkacb

### Cholinergic synapse in STR-CP&ACB

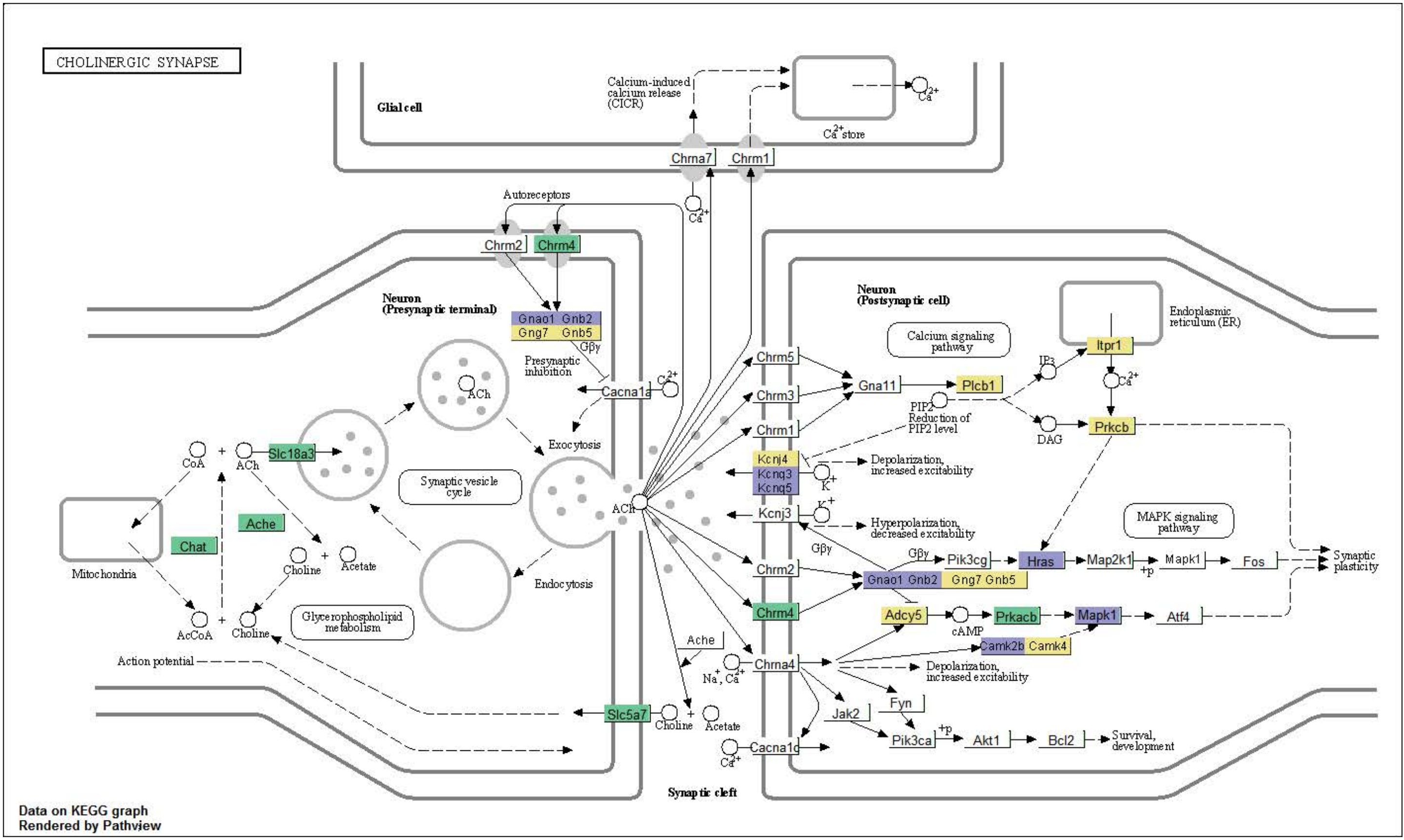


**Pre:** Gnao1 Gnb2
**Pre:** Gng7 Gnb5
**Pre:** Chat Ache Slc18a3 Slc5a7 Chrm4

**Post:** Kcnq3 Kcnq5 Gnao1 Gnb2 Camk2b Hras Mapk1
**Post:** Kcnj4 Adcy5 Plcb1 Itpr1 Prkcb Camk4 Gng7 Gnb5
**Post:** Chrm4 Prkacb

#### GABAergic synapse in STR-CP&ACB

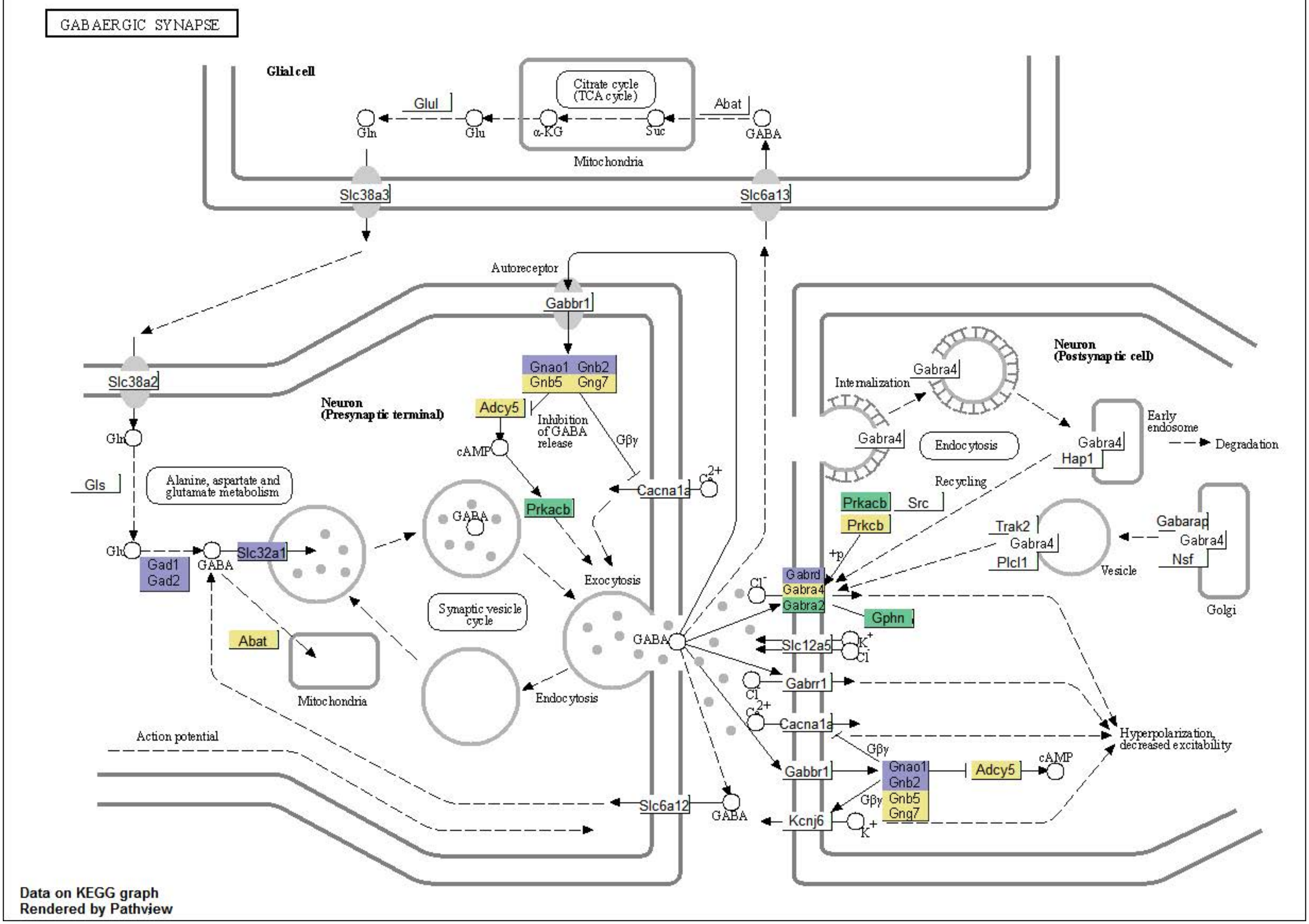


**Pre:** Gad1 Gad2 Slc32a1 Gnao1 Gnb2
**Pre:** Abat Adcy5 Gnb5 Gng7 Gabra4 Prkcb
**Pre:** Prkacb

**Post:** Gnao1 Gnb2 Gabrd
**Post:** Adcy5 Gnb5 Gng7 Gabra4 Prkcb
**Post:** Gphn Gabra2 Prkacb

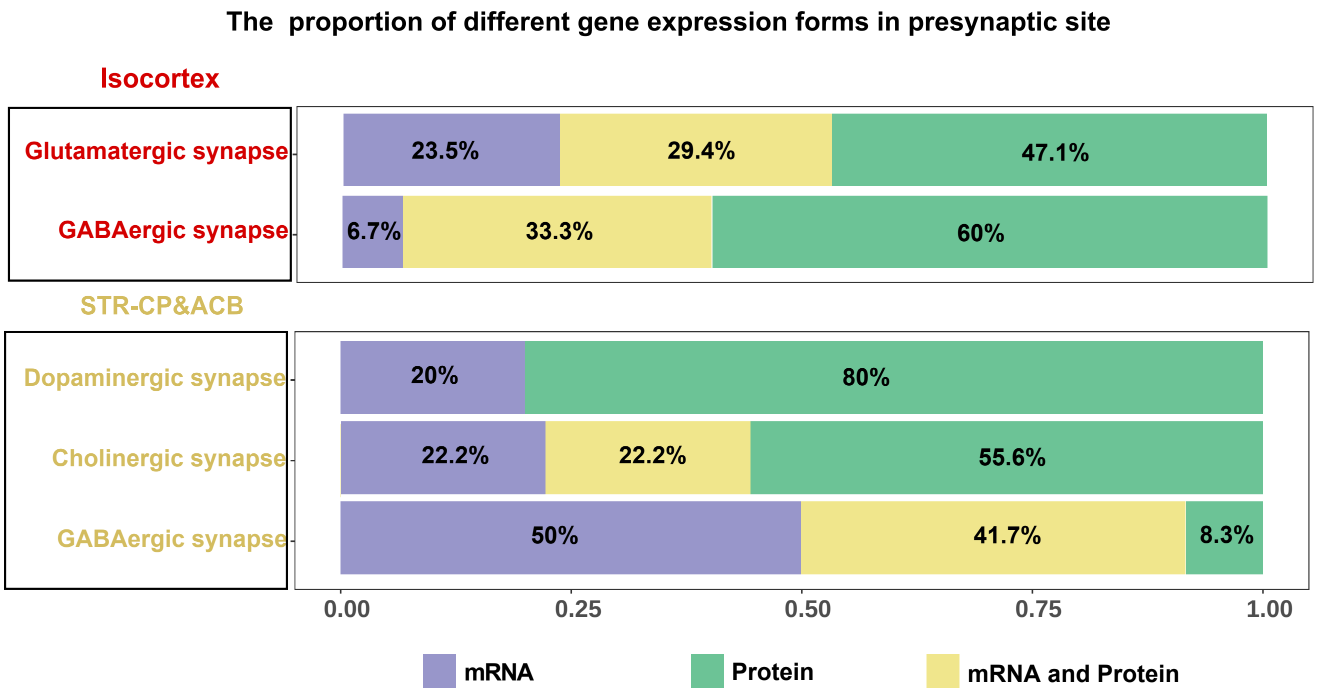

The proportion of different gene expression forms in presynaptic site
Isocortex
Glutamatergic synapse
GABAergic synapse
23.5%
29.4%
47.1%
6.7%
33.3%
60%
STR-CP&ACB
Dopaminergic synapse
Cholinergic synapse
GABAergic synapse
20%
80%
22.2%
22.2%
55.6%
50%
41.7%
8.3%
0.00
0.25
0.50
0.75
1.00
mRNA
Protein
mRNA and Protein
