## Supplementary figures and images for "Mouse Forebrain Region-specific Molecular Characteristics and Cholinergic Neurons Subtyping: An Integrated Analysis Based on Spatial Multi-omics"

### Figure S2.pdf

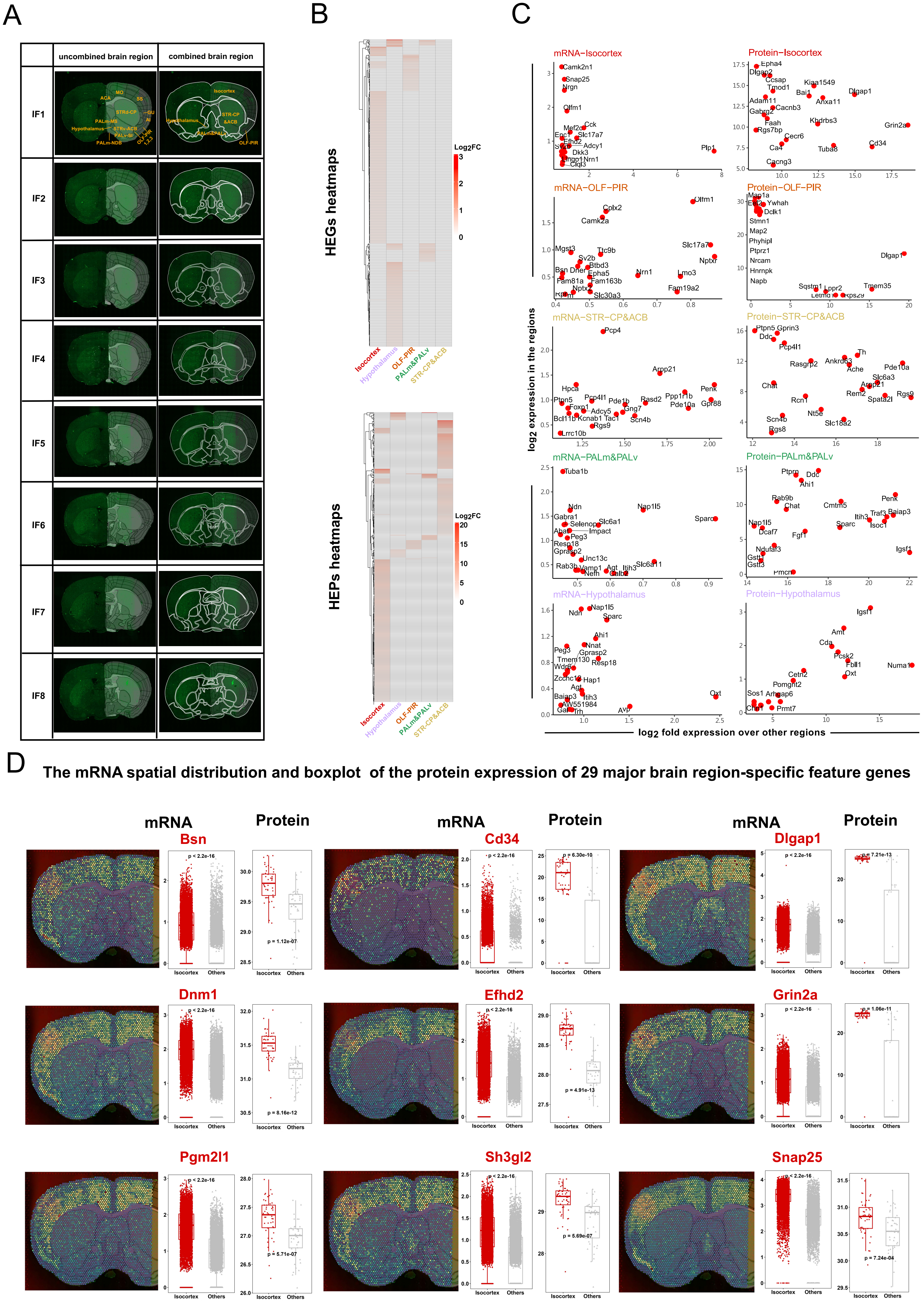

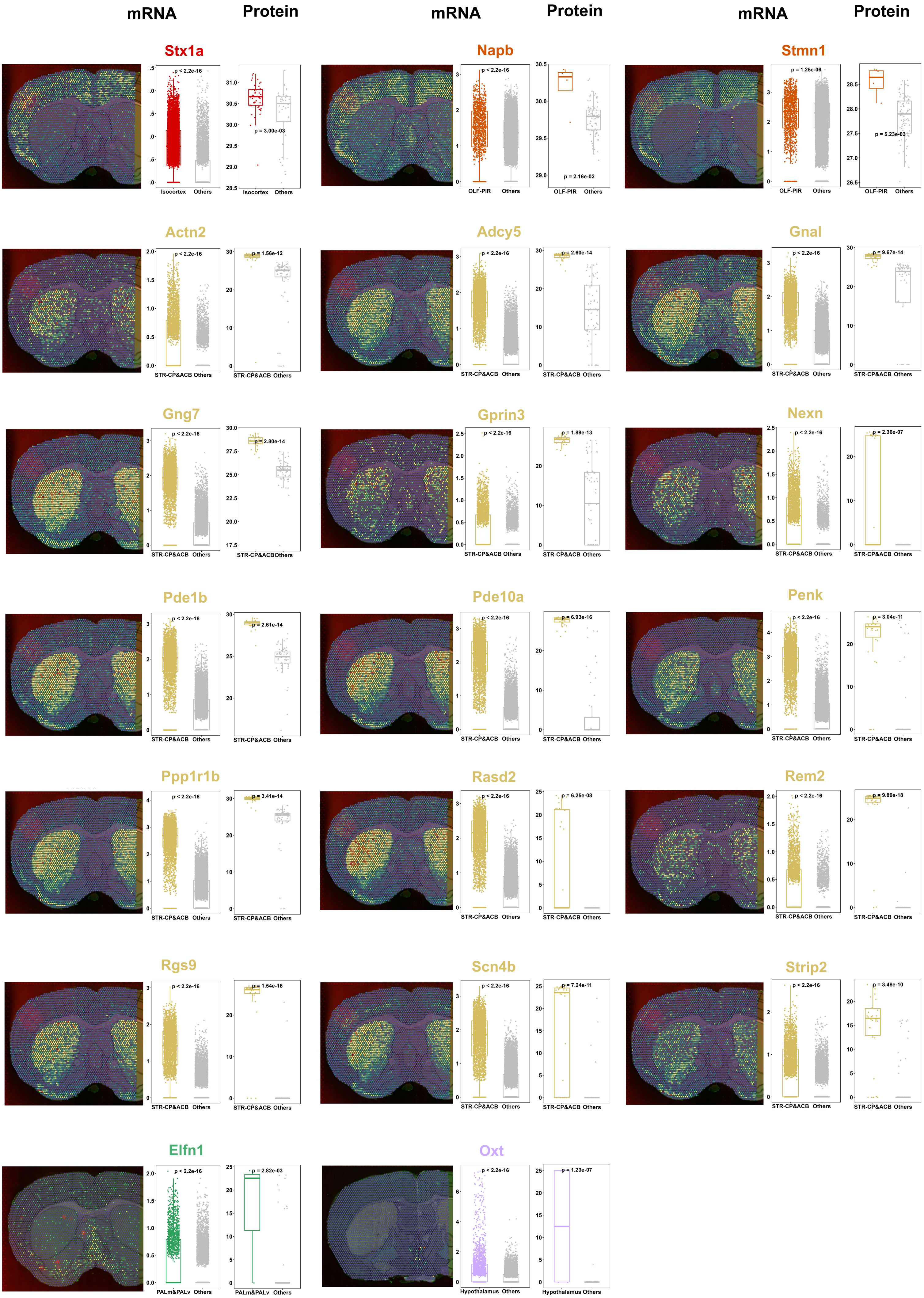

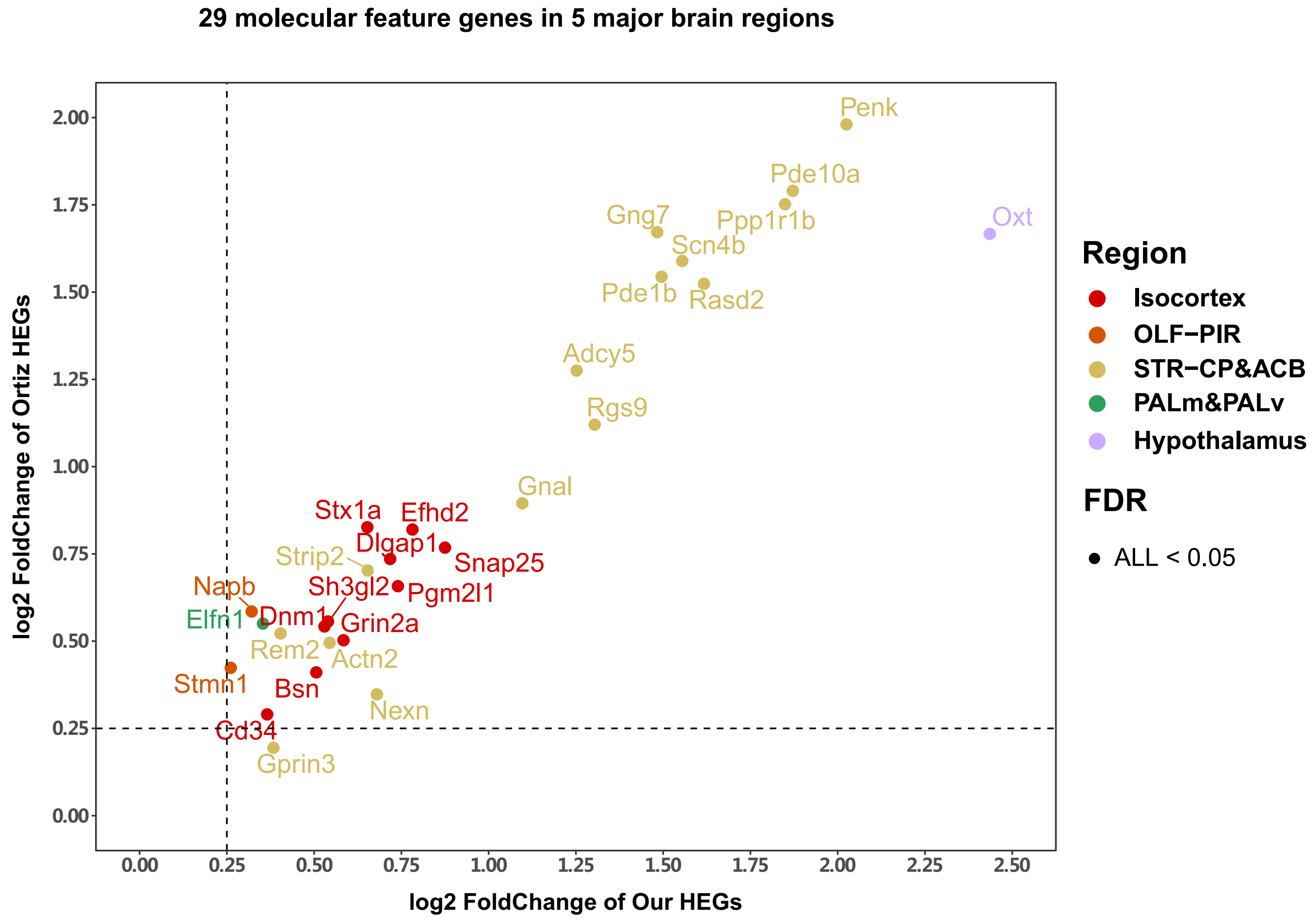

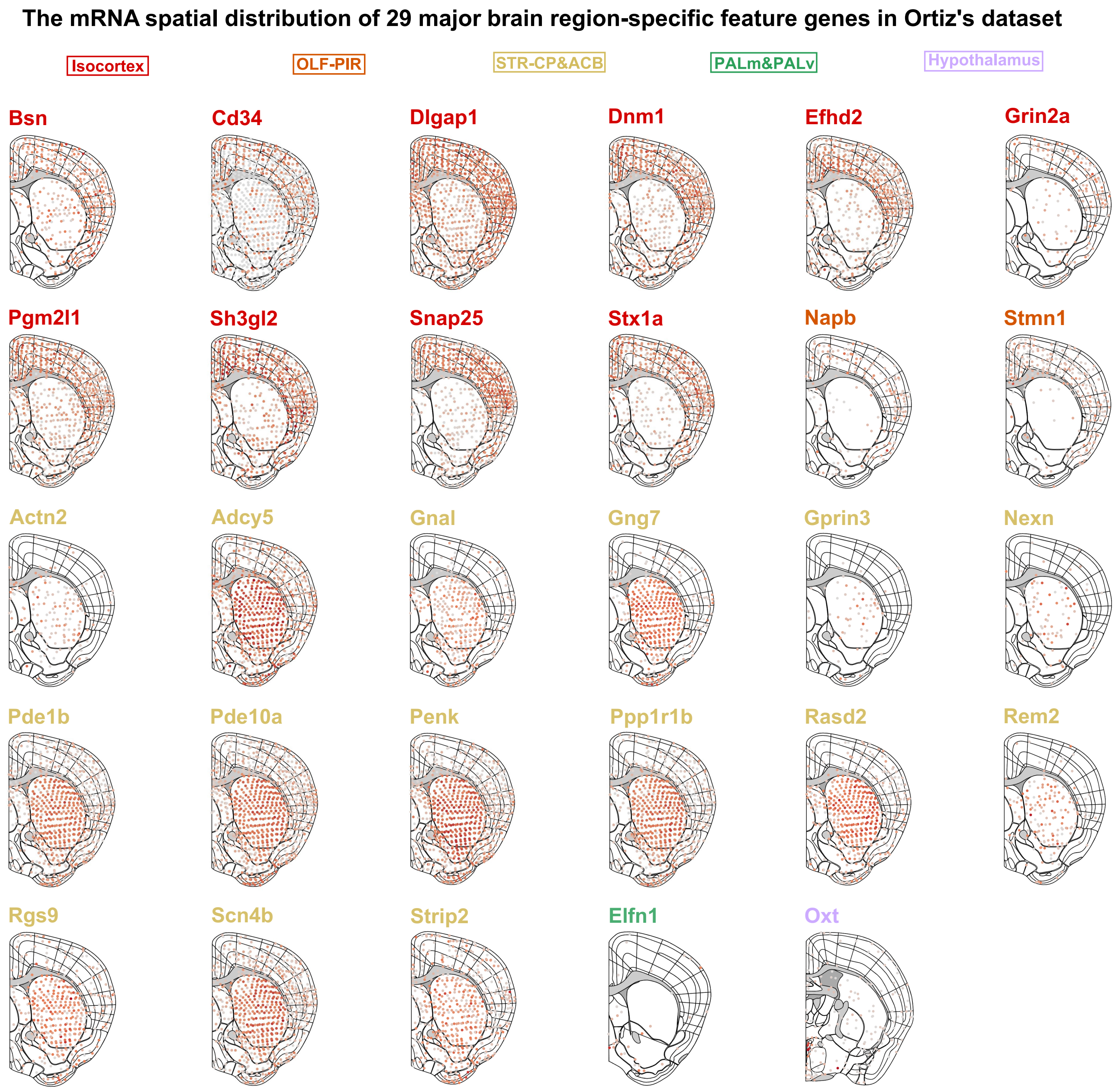

### Figure S3.pdf

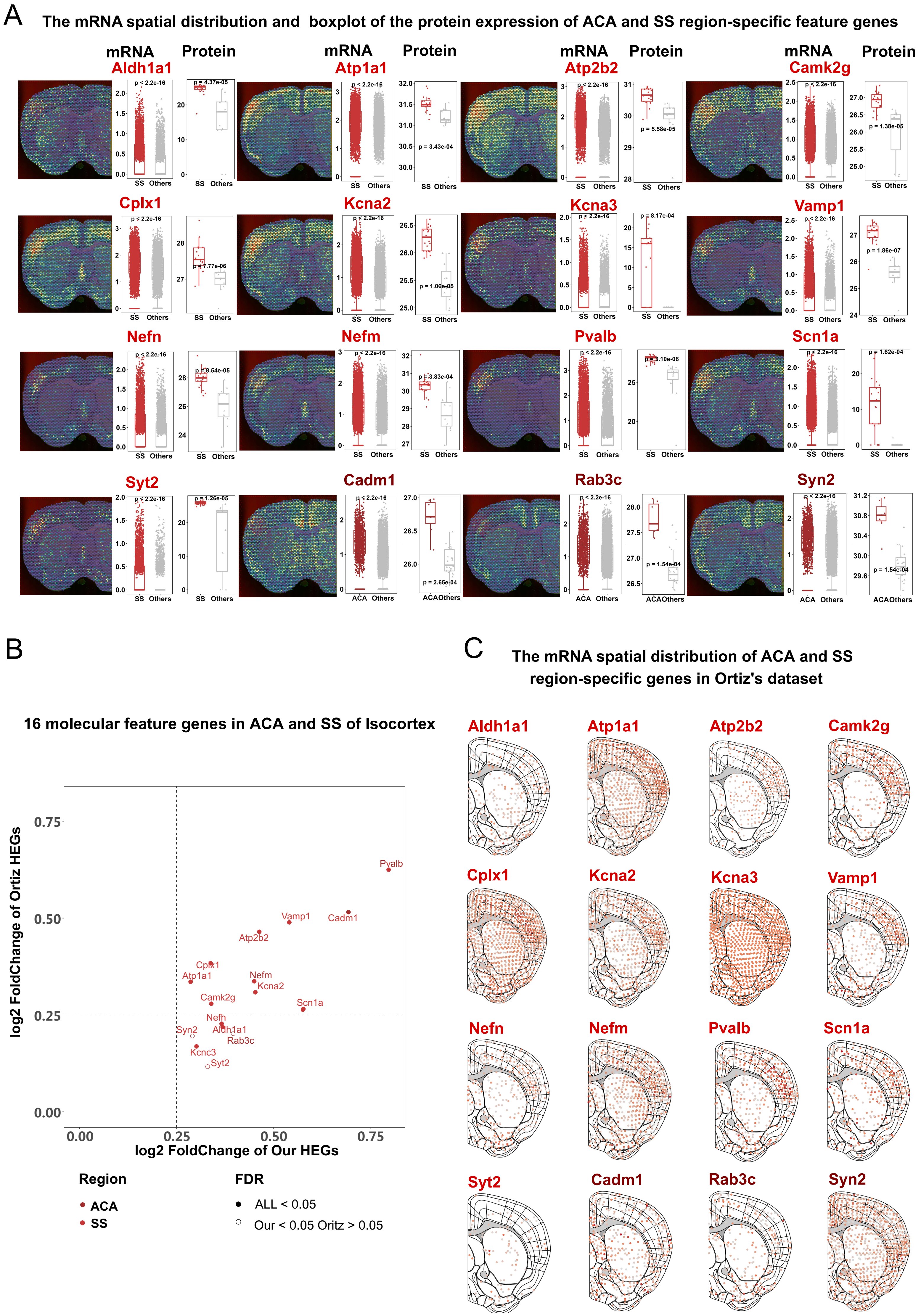

### Figure S6.pdf

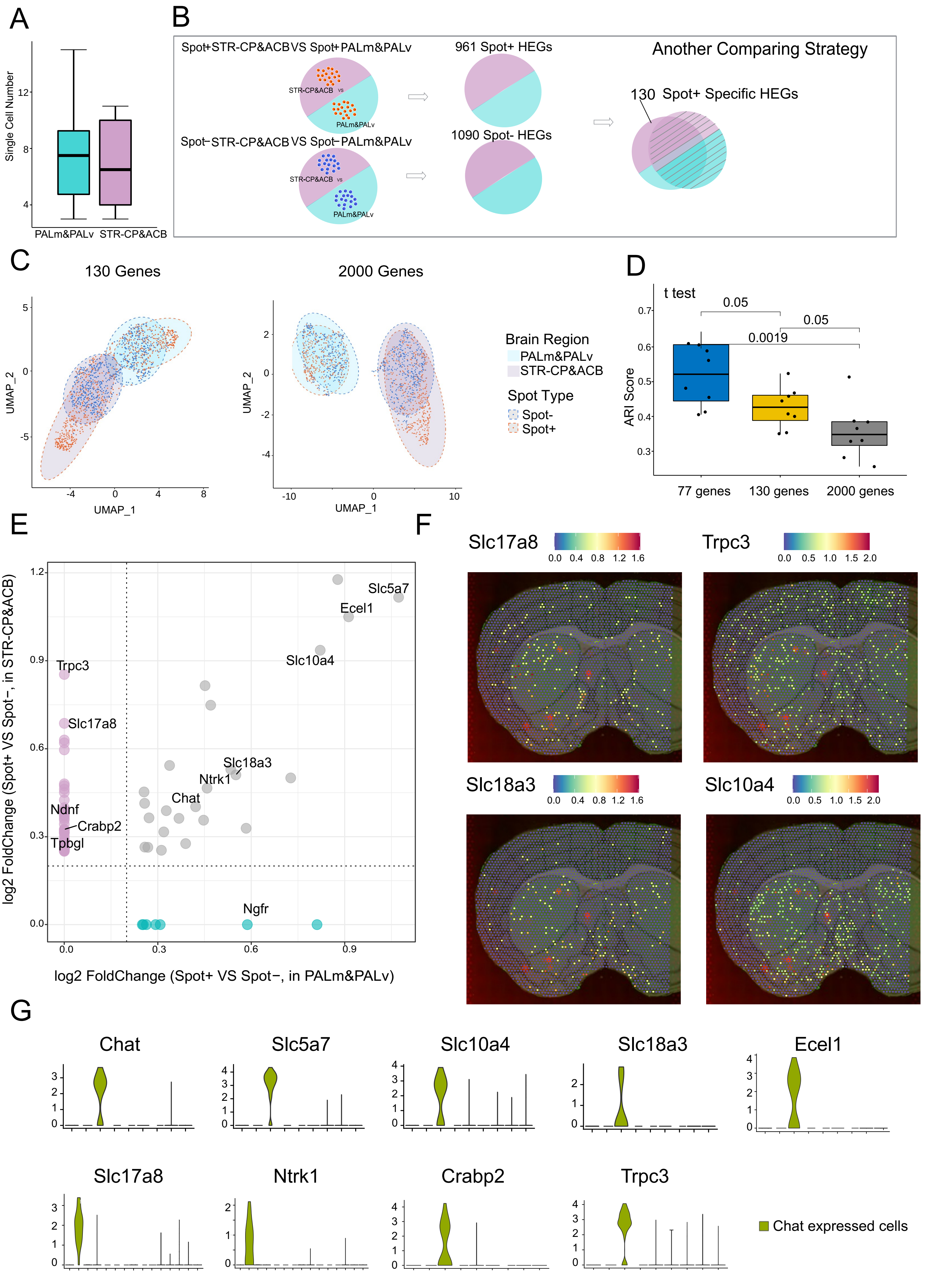

### Figure S7.pdf

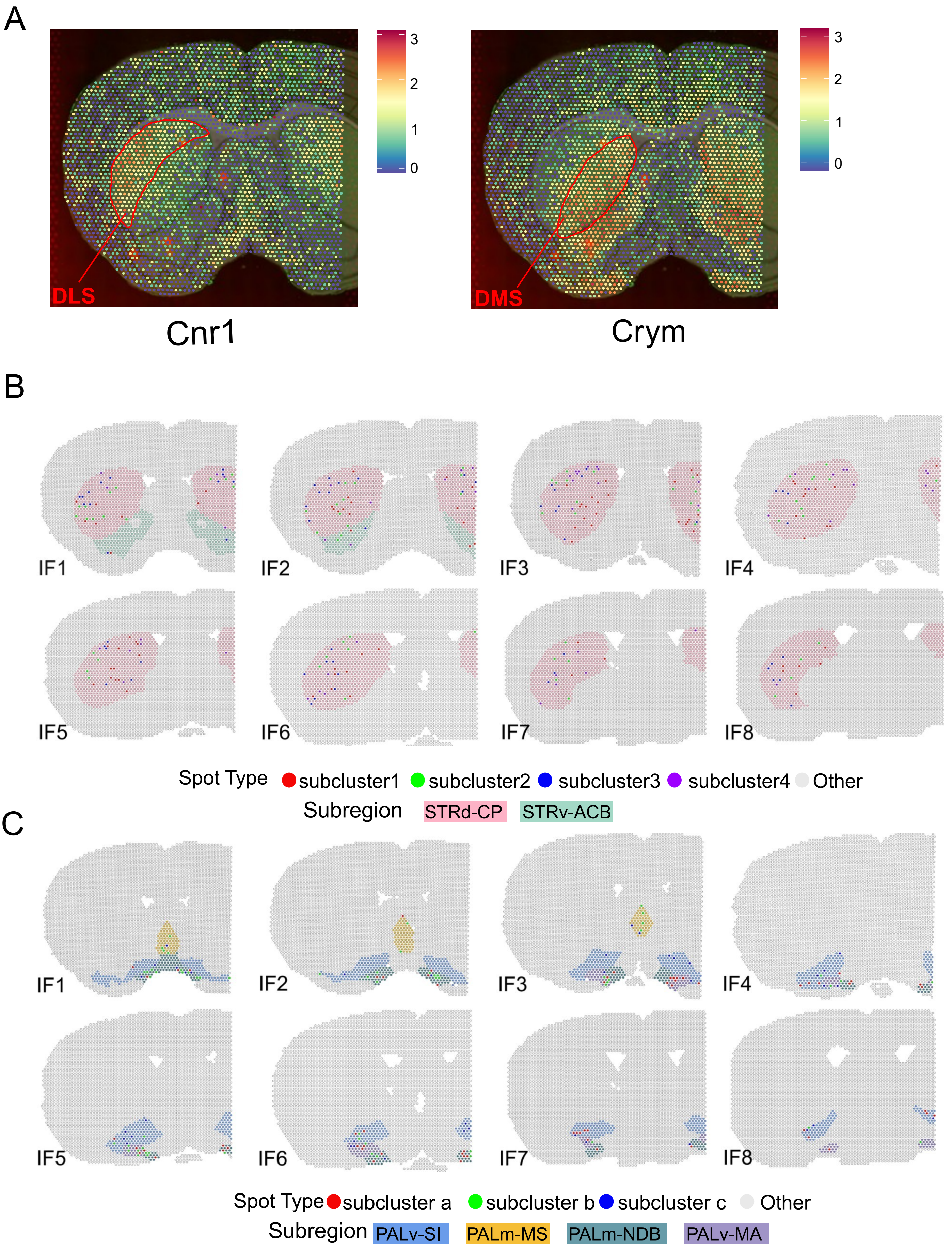

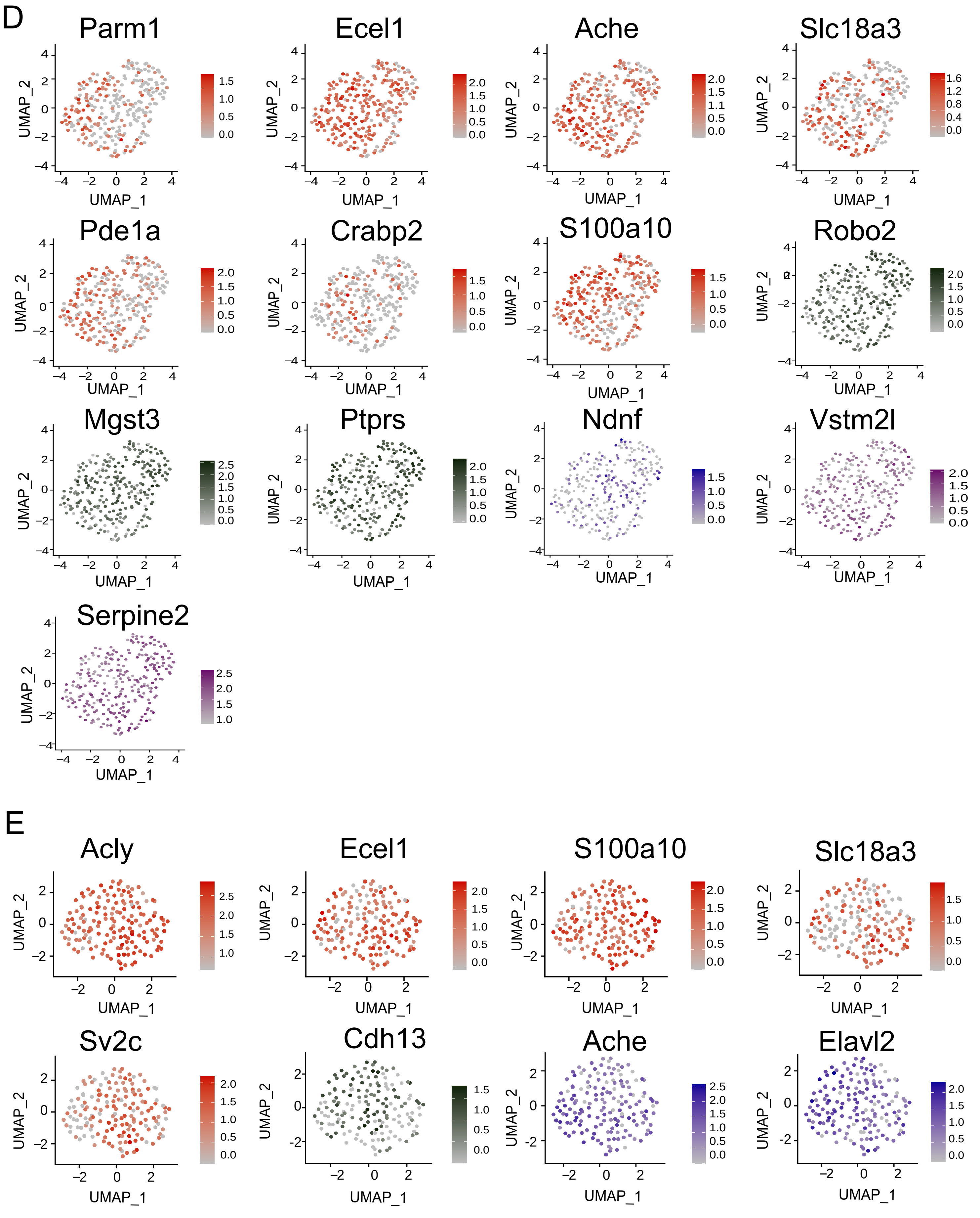

### Figure S8.pdf

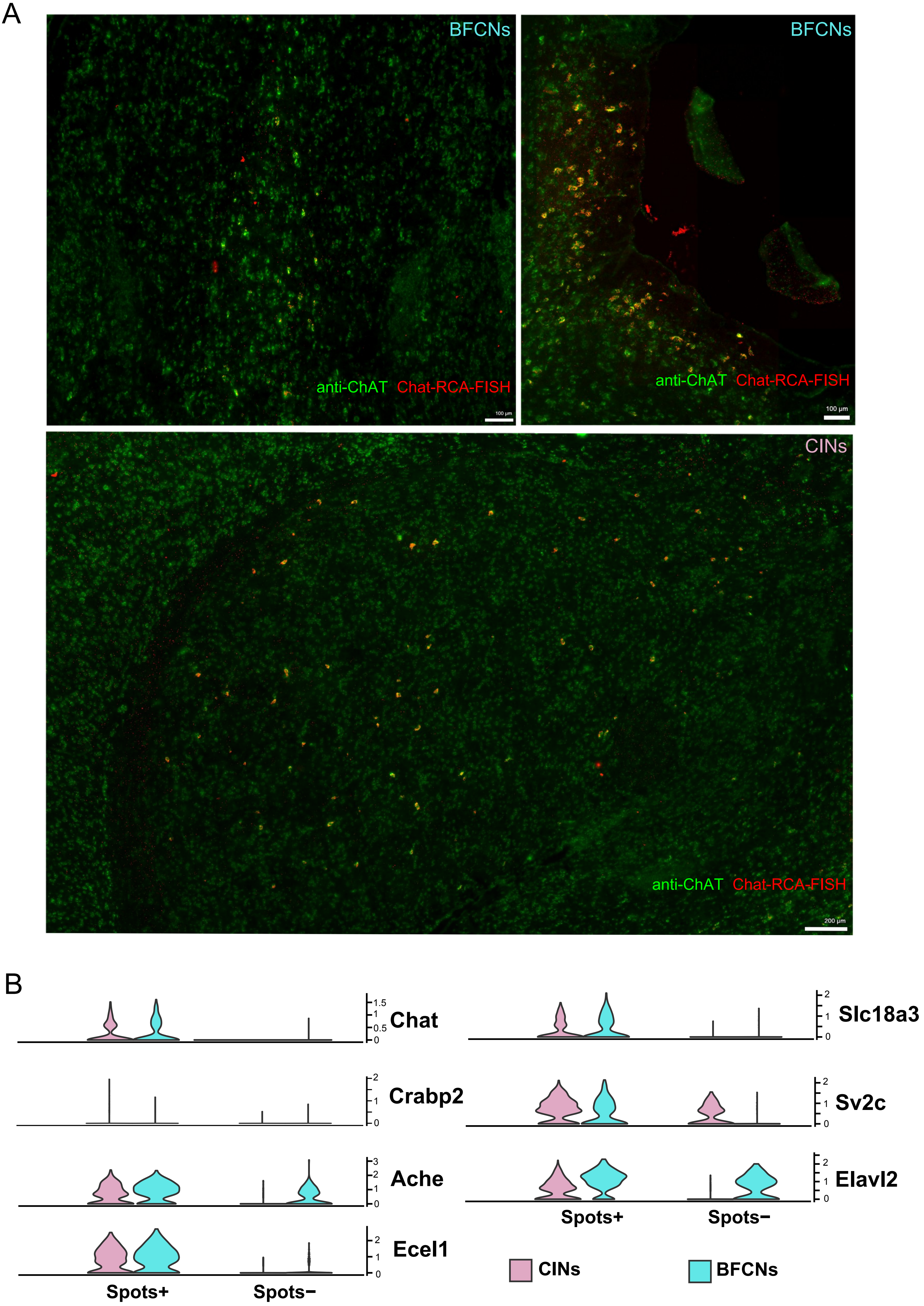

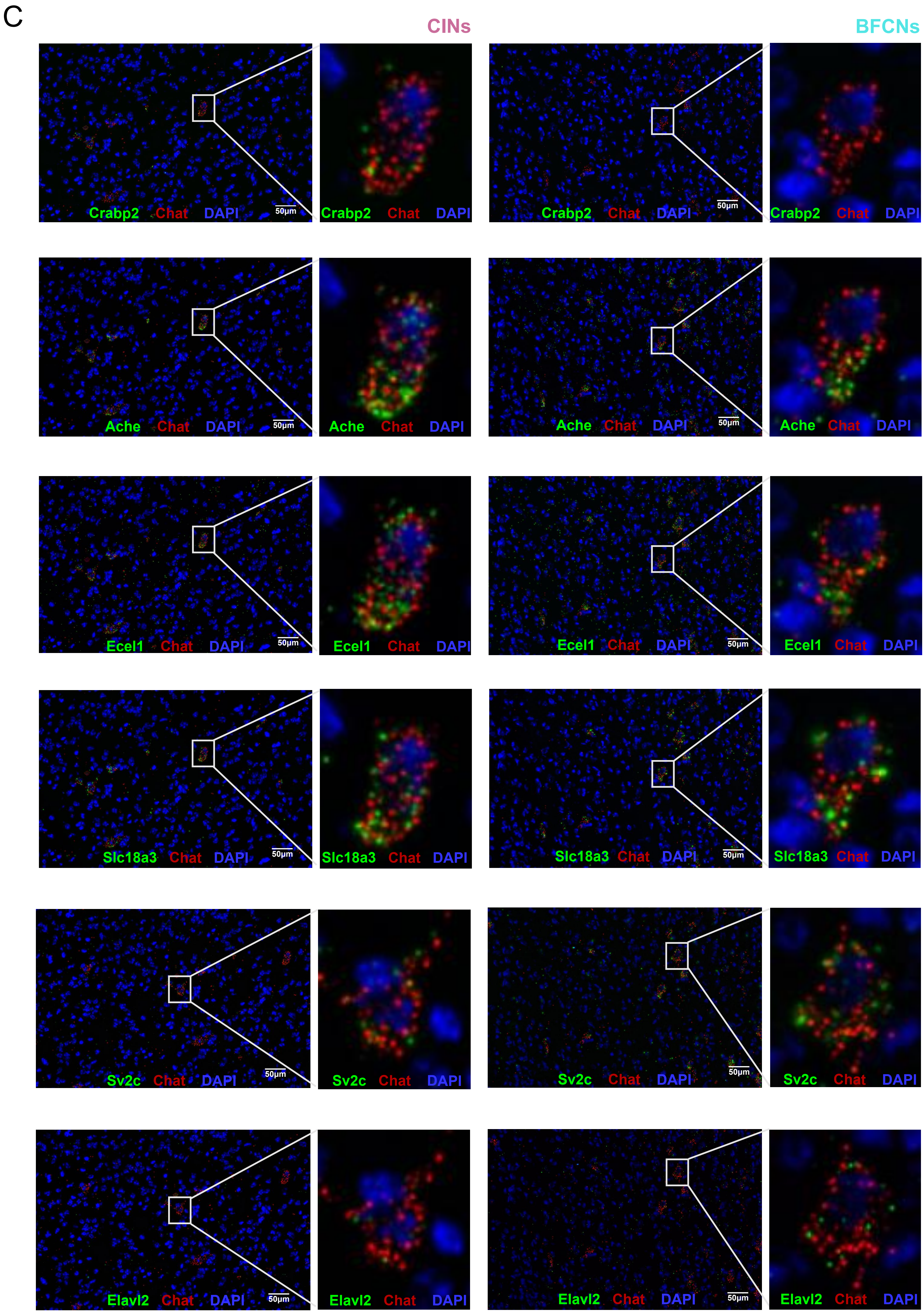
